# Analysis of the Hamiltonian Monte Carlo genotyping algorithm on PROVEDIt mixtures including a novel precision benchmark

**DOI:** 10.1101/2022.08.28.505600

**Authors:** Mateusz Susik, Ivo F. Sbalzarini

## Abstract

We provide an internal validation study of a recently published precise DNA mixture algorithm based on Hamiltonian Monte Carlo sampling [1]. We provide results for all 428 mixtures analysed by Riman et al. [2] and compare the results with two state-of-the-art software products: STRmix™ v2.6 and Euroformix v3.4.0. The comparison shows that the Hamiltonian Monte Carlo method provides reliable values of likelihood ratios (LRs) close to the other methods. We further propose a novel large-scale precision benchmark and quantify the precision of the Hamiltonian Monte Carlo method, indicating its improvements over existing solutions. Finally, we analyse the influence of the factors discussed by Buckleton et al. [3].

## 1 Introduction

Robust interpretation of mixed DNA profiles is at the core of forensic practice. Algorithmic and computational support for this task has greatly evolved over time, and the most recent software generation of *fully continuous* methods can accurately resolve traces that were previously considered as unresolvable [4]. Given that every mistake of such algorithms could have drastic consequences (e.g. providing false evidence against a suspect), any software in practical use has to be thoroughly validated and trusted by the forensic community. Laboratories intending to adopt new technology should therefore have a way to assess the correctness and reproducibility of the results produced by a given genotyping software. A common way of validation is to check the results of the tested algorithms on a set of known DNA mixtures prepared under laboratory conditions [2, 5, 6]. Statistical analysis of the results can reveal the discriminatory power of the algorithms.

All laboratories are therefore recommended to validate the software they use before applying it in an investigation process [7, 8]. The effort required to adopt fully continuous methods is generally considered higher than with older techniques, especially for software products that recommend estimation of laboratory-dependent hyper-parameters. Since most potential users do not have the possibility of preparing their own in-house validation mixtures, it should be possible to compare them without the need to perform a laboratory pipeline.

This requirement led to increasing efforts towards comparative evaluations of different methods, resulting in benchmark studies that offer convenient summaries of the discriminatory power of the available products [2, 5]. These studies are performed on standardized mixtures from the openly available PROVEDIt (Project Research Openness for Validation with Empirical Data) dataset [9], and thanks to the study by Riman et al. [2] it became possible to validate new inference methods by comparing them with previous solutions. Another reason for the importance of comparative studies is that they offer a possibility to estimate the effects of differences between solutions on the output likelihood ratios (LRs). This reason has been highlighted in the President’s Council of Advisors on Science and Technology (PCAST) report [8].

Most comparative studies so far have focused on two software solutions: Euroformix [10] and STRmix™ [11]. The latter uses Random-Walk Markov Chain Monte Carlo (MCMC) to estimate the probabilities of possible genotype sets. The former was benchmarked using its maximum-likelihood estimations (MLE) method. Euroformix also offers an MCMC sampler (referred to as “Bayesian method” in that software), which, however, has never been considered in comparative benchmarks.

The results of MCMC techniques are stochastic approximations, and the output LRs are expected to exhibit run-to-run variability. The magnitude of this variability strongly depends on the design of the probabilistic genotyping model and on the sampling algorithm used, and it can be significantly reduced by appropriate choices [1]. This was demonstrated by using a Hamiltonian Monte Carlo (HMC) sampling algorithm, which leverages Hamiltonian dynamics over the optimization landscape in order to propose new samples that have high acceptance probability and are uncorrelated to past samples. This requires computing the gradient of the unnormalized log-posterior, which can be efficiently done using automatic differentiation frameworks. For maximum-likelihood estimators (MLE), variability is not expected. However, it might still be observed in practice if the algorithm is not able to find the global maximum in all runs or if the convergence was not properly assessed.

So far, run-to-run variability is rarely systematically benchmarked, and no comparative evaluation of HMC-based methods exists in the literature. Here, we therefore validate the HMC-based probabilistic genotyping method of Susik et al. [1] on the publicly available PROVEDIt mixtures [9], and we propose a novel precision benchmark for quantifying run-to-run variability. We compare the results with those provided by STRmix™ v2.6 [2] and Euroformix v3.4.0. Doing so, we provide additional arguments for the robustness and correctness of the results obtained by the HMC method. We also provide a wider perspective for comparing of different software solutions. Given that the probabilistic model of the tested HMC method is shared with one of the comparison methods [11], the present results also shed light on the reproducibility of published studies. By comparing the results of the HMC method with those obtained by STRmix™ and Euroformix, we quantify the influence of the choice of the probabilistic model and of software implementation differences. We specifically analyse the effects of the choices mentioned by Buckleton et al. [3] when discussing the results of Riman et al. [2]. We hope that the results presented here foster quantitative comparisons and methodologically founded discussions about the precision of the available algorithms and the reproducibility of their results.

## 2 Methods

We perform an internal validation study on publicly available mixtures that were also used in previous comparative studies [2]. This allows us to compare the results between studies. Riman et al. [2] reported results for one MCMC implementation of a log-normal model (STRmix™) and one MLE implementation of a gamma model (Euroformix). The HMC method considered in this study uses a log-normal model closely following STRmix™’s methodology.

### 2.1 PROVEDIt mixtures

The PROVEDIt database contains mixtures of varying contributors, numbers of contributors, contributor ratios, and mixture qualities [9]. In many cases the mixtures have been deteriorated by treatments with DNAse I, Fragmentase^®^, UV irradiation, sonification, or humic acid. The database contains mixtures analysed with different multiplexes, instrument models, and injection times.

We follow the study conducted by NIST [2] and perform our analysis on mixtures amplified with the GlobalFiler kit with 29 PCR cycles and analysed on a 3500 Genetic Analyzer with an injection time of 15 seconds. The filtered input files are used.

### 2.2 Past discussion on published Euroformix results

Buckleton et al. [3] criticise the choices made by Riman et al. [2]. The critique is summarised in a list of 5 points. In order to be able to compare HMC with the only available larger STRmix™ benchmark, we follow the study of Riman et al., but take the following measures:

- We re-analyse 2- and 3-contributor mixtures with Euroformix 3.4.0 with a set of hyperparameters we estimated. This responds to points 1 and 3 raised by Buckleton et al. [3].
- We compare different rare-allele models based on results obtained from the same likelihood ratio system [2]. This responds to point 4 raised by Buckleton et al. [3].

Point 2 raised by Buckleton et al. [3] criticises the use of a custom tool to pre-filter artefact peaks. It has been independently verified (see Supplementary Material of Ref. [1]) that this tool has indeed filtered out some of the stutter peaks that are important for the analysis, so this is definitely an issue. However, the study of Riman et al. [2] is the only paper available that makes it possible to compare with STRmix™ results, which is why we nevertheless use it.

It is our aim to provide a fair comparison. Still, as we have to estimate the hyperparameters for Euroformix based on different modelling choices (see Subsection 2.3), the comparison is not perfect. We nevertheless choose this way because Euroformix v3.4.0 does not model double-backward stutter. An alternative approach would be to force the same limitation onto the more versatile algorithms, similarly to how it has been done by Cheng et al. [5]. However, laboratories that adopt solutions modelling more types of stutters will intend to use those features (e.g. to permit lower analytical thresholds). Therefore, we feel that such a comparison would be of limited relevance to practical casework.

### 2.3 Hyperparameters settings

In order to reduce possible sources of differences between inference methods, we reuse the parameters calculated for STRmix™ in Ref. [2] with the exception of the parameters for the drop-in model. Since HMC uses the drop-in model from Euroformix, we reuse the published hyper-parameters from Euroformix for the drop-in model. The values of all hyperparameters are available in the original paper on the HMC algorithm [1]. Unless specified otherwise, we report sub-source LRs, because they do not reflect any assumption about the contributor type of the person of interest (e.g. major vs. minor). We resort to sub-sub-source LRs whenever we analyse a mixture in depth, for example when we report an LR for an individual locus.

Euroformix v3.4.0 enables modelling of forward stutters in addition to backward stutters. Still, no model for double-backward stutters is available yet. To follow these assumptions without biasing the results, we use higher analytical thresholds (ATs), such that double-backward stutters are filtered out and will not distort the output. We determine these ATs by analysing the available single-source profiles (Supplementary Material 1) and removing the peaks of the contributors, backward, and forward stutters. We follow Riman et al. [2] and use the formula AT = *µ* + 10*σ*, where *µ* and *σ*are the mean and standard deviation estimated from the remaining peak heights (including double-backward and half-backward stutters). The resulting ATs used for LR calculation are: 60, 80, 45, 75, and 100 RFU for the blue, green, yellow, red, and purple dyes, respectively. Under these ATs, neither double-backward nor half-backward stutters were left in the 2- and 3-contributor mixtures. We confirmed this by comparing the left peaks with the ground-truth contributors’ genotypes. In practical casework, however, this information is not available to the laboratory, and they need to find another way to deal with the issue. Our higher ATs result in analysing 2729 peaks less than Riman et al., namely an average of 6.38 per a mixture. Degraded mixtures are affected more, given the lower heights of their allelic peaks on the right side of the dyes.

A single drop-in model is estimated from the same single-source profiles as the ones used for AT estimation. The estimated drop-in probability is 0.00073 and the lambda is 0.03846. An *F*_*ST*_ correction of 0.01 is used, and allele frequencies are normalised.

For Euroformix, we only provide results for 2- and 3-contributor mixtures. This is for reasons of analysis runtime, which were too large (regularly exceeding an hour per scenario) to be practical for the required numbers of repetitions when using Euroformix on 4-contributor mixtures. There are at least two reasons for this: (1) In order to provide results for different PoIs (persons of interest) under the same hypothesis of the prosecutor, the inference process has to be run separately for each PoI in Euroformix. In the case of HMC, the inference can be performed once, as the LRs for different PoIs can re-use the same estimation of the posterior probability distribution. (2) In cases with few or no stutter peaks, the stutter model used by Euroformix might not converge. The inference might then never provide an answer. In such cases where the Euroformix model does not converge, we switch off the stutter model that caused the issue (firstly forward, and then if it did not help, also the backward model). Such a strategy causes an interesting situation in which we report results with different settings of stutter models within the same mixture. Still, we believe this is the approach that would imitate the best an approach adopted in a laboratory, as often the laboratories will test only one hypothesis of prosecutor. The list of the resulting stutter model settings is available in Supplementary Material 1.

### 2.4 Benchmark framework

All compared software products compute likelihood ratios [12]. This ratio expresses the strength of the evidence and is defined as:

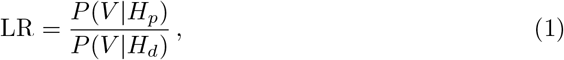

where *V* is the observed electropherogram (EPG), and *H*_*p*_ and *H*_*d*_ are the hypotheses of the prosecutor and the defendant, respectively. Each *H*_*p*_ in our benchmark assumes presence of the PoI. All hypotheses in the benchmark assume that the contributors are of Caucasian ethnicity and come from the U.S. population genetic background. Therefore, NIST 1036-Caucasian allele frequencies based on 361 individuals are used. The hypotheses include the information about the number of contributors. In all cases, *H*_*p*_ assumes the inclusion of a certain person of interest (PoI), while *H*_*d*_ assumes that all contributors are unknown. For each mixture, all possible scenarios with a true contributor in *H*_*p*_ are considered against the same number of scenarios with false PoIs (i.e. non-contributors). All contributors are assumed to be unrelated. For example, a contributor mixture will be analysed with 6 different hypotheses: 3 with a true contributor and 3 with a false one. The false PoIs has been chosen randomly from the genotypes of the NIST 1036-Caucasian sample [2]. The benchmark consists of 154 two-contributor mixtures, 147 three-contributor mixtures, and 127 four-contributor mixtures. We define:

- Scenario: A scenario is a setting where a mixture, *H*_*p*_, and *H*_*d*_ are defined. For each mixture in the benchmarks, multiple scenarios are provided.
- Person of interest (PoI): The PoI is the person (and his/her genotype) assumed present in the mixture under the prosecutor’s hypothesis, but not under the defendant’s hypothesis.
- True/false hypothesis: An *H*_*p*_ in which the PoI is a contributor to the sample is called a true hypothesis. If the PoI is a non-contributor, then *H*_*p*_ is called a false hypothesis.
- Opposite of the neutral threshold (OotNT): A scenario is opposite of the neutral threshold (LR = 1) if the software outputs LR*>*1 for a false hypothesis or LR*<*1 for a true hypothesis.
- OotNT rate: The proportion of scenarios that are OotNT for a given probabilistic genotyping software.
- Precision: Precision measures reproducibility of the results when the same analysis run is repeated multiple times. The lower the run-to-run variation, the more precise the analysis. Precision depends on the scenario, hyperparameter values, background population frequencies and is quantified by the LR variability.
- Ban: A ban is a logarithmic unit for base 10 logarithms. 1 ban = 10, 2 bans = 100, 3 bans = 1000, etc.

## 3 Results

We first analyse the classification performance of the HMC method in comparison with the established approaches. Then, we introduce a novel large-scale benchmark for quantifying and comparing the precision of the different methods. Finally, we quantify differences that result from different rare-allele models used by the different methods.

### 3.1 Classification performance

Probabilistic genotyping systems are required to rarely provide evidence in favour of an untrue hypothesis for PoIs that are unrelated to the true contributors. Indeed, Slooten [13] noted:

> “To report evidence in favour of another hypothesis than the one that is actually true is not an error when the data have been correctly analysed for the relevant hypotheses, because then the data were simply misleading, but one could consider it an error, or at least very undesirable, when it is reported that there is evidence whereas there is none, due to an inadequate model having been used.”

Therefore, OotNT scenarios are not necessarily a sign of malfunctioning software. However, large OotNT rates do indicate that the software either does not follow the principles outlined by Slooten, that the model is unable to explain the peak heights, or that the software code contains programming mistakes. We cannot know which of the OotNT scenarios would be still OotNT if we had Slooten’s hypothetical “perfect” probabilistic genotyping system. The neutral threshold we choose (LR=1) has the natural statistical meaning of delineating two alternative hypothesis. However, testing other thresholds, for example higher thresholds for multiple-hypotheses testing in database searches, is also useful.

We provide a comparison of OotNT cases, showing that HMC achieves state-of-the-art results. Indeed, we find that HMC is characterised by a low OotNT rate, comparable to that of STRmix™ and lower than that of Euroformix (Table 1). For the 4-contributor mixtures the OotNT rates of the MCMC-based methods are under 2.5% (2.1% for HMC and 2.4% for STRmix™), while Euroformix provided 16.1% OotNT scenarios in the original study [2]. For Euroformix, the hypotheses with non-contributors dominate the OotNT cases with a total of 70 versus only 2 true-contributor OotNT scenarios across all 2- and 3-contributor mixtures. This is in contrast to the MCMC-based methods more often providing OotNT LRs for true hypotheses (for 21 true vs. 13 false hypotheses in the case of HMC, 26 true vs. 17 false in the case of STRmix™).

**Table 1.** OotNT rates and absolute numbers of OotNT scenarios with true contributors and non-contributors. We compare three analysis methods in scenarios with different numbers of contributors (NoC).

| NoC | HMC |  |  | STRmix<sup>TM</sup> |  |  | Euroformix |  |  |
| --- | --- | --- | --- | --- | --- | --- | --- | --- | --- |
|  | Rate | Non-contr. | Contr. | Rate | Non-contr. | Contr. | Rate | Non-contr. | Contr. |
| 2 | 0.002 | 0 | 1 | 0.008 | 1 | 4 | 0.023 | 13 | 1 |
| 3 | 0.014 | 2 | 10 | 0.016 | 2 | 12 | 0.066 | 57 | 1 |
| 4 | 0.021 | 11 | 10 | 0.024 | 14 | 10 | — | — | — |

We further analyse the OotNT scenarios in Figure 1. Most of them do not provide strong evidence, with the majority of log_10_ LR between -1 and 1: 67.6% (23/34) for HMC, 69.8% (30/43) for STRmix™, and 93.1% (67/72) for Euroformix.

**Figure 1.**
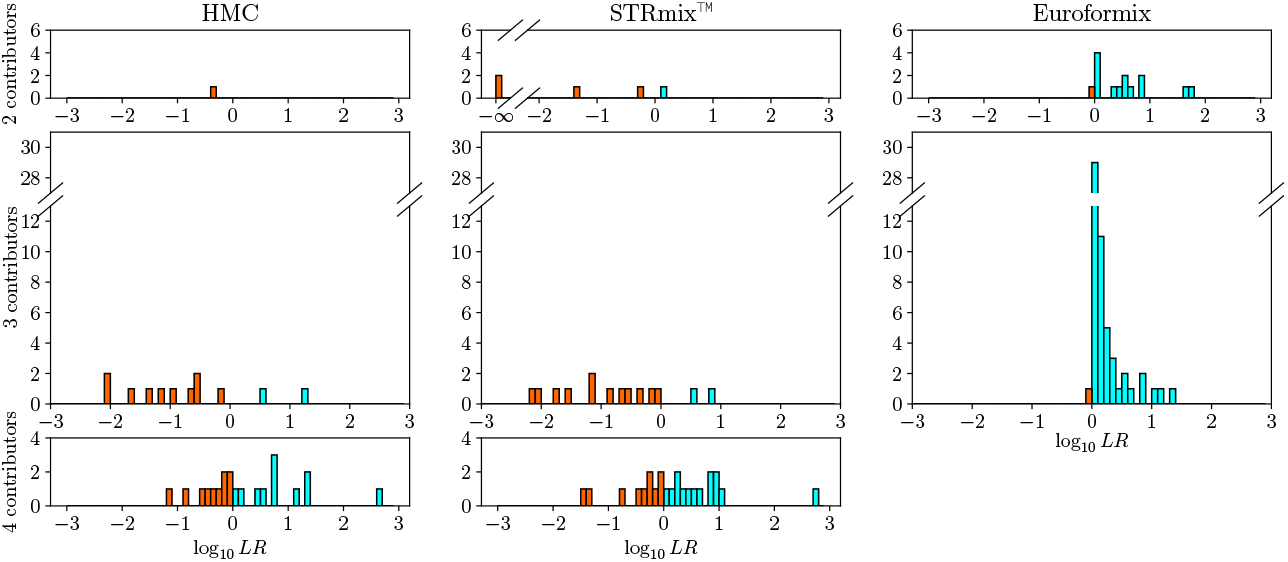
Histogram of OotNT scenarios. Cyan represents OotNT scenarios with false contributors and orange the ones with true contributors.

Another popular metric for evaluating the quality of a classification method is the area under the Receiver Operating Characteristic (ROC) [14]. We therefore plot the ROC curves for the tested methods in Figure 2. All tested methods achieve performances with normalised areas under the curves close to the maximum theoretical value of 1.0^1^, see inset legends. The slight difference between STRmix™ and HMC is mainly caused by two OotNT scenarios with LR=0 from STRmix™. Increasing the number of STRmix™ iterations in these scenarios increases the area under the ROC curve for STRmix™ from 0.99600 to 0.99994 for the 2-contributor mixtures. This is then perfectly comparable to the performance achieved by HMC.

**Figure 2.**
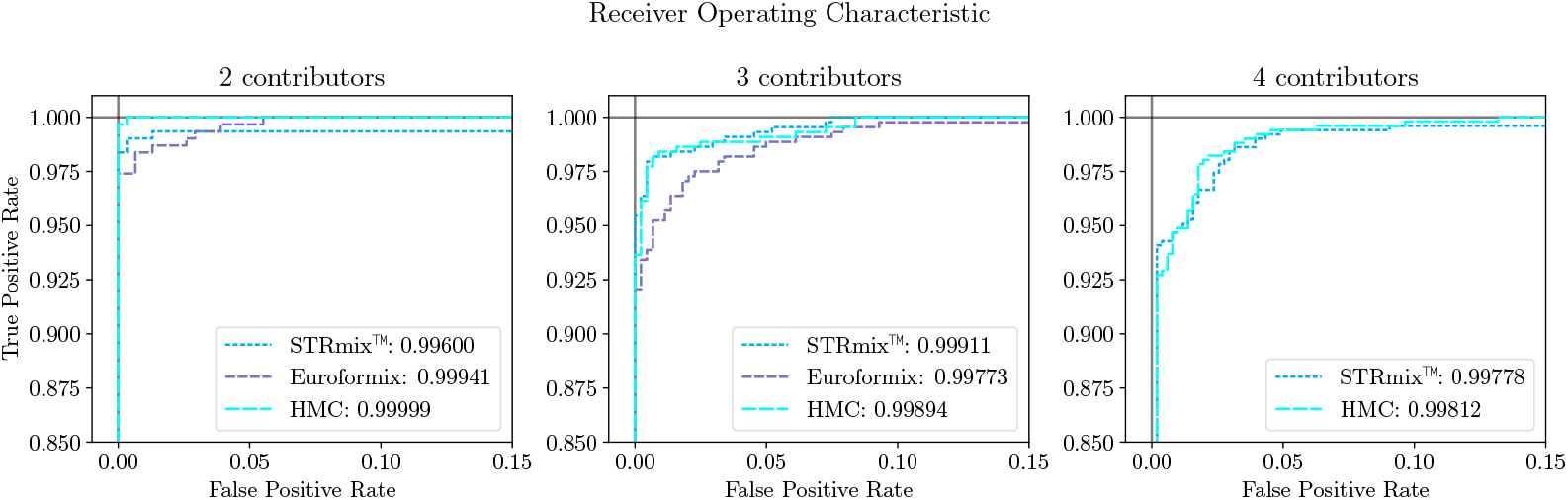
Comparison of Receiver Operating Characteristic (ROC) curves between the tested methods. The normalized areas under the ROC curves are given in the inset legends. They measure classification performance with a perfect classifier achieving 1.0.

The two false negatives responsible for this behaviour are two of the true hypotheses among the 2-contributor mixtures for which STRmix™ falsely outputs LR=0. Riman et al. [2] therefore decided to re-run the analysis on these cases with a larger number of iterations. Those repeated runs provided strong evidence for the inclusion of the respective PoIs (log_10_ LR of 24.88 for the mixture marked as C02 and 19.66 for case H06). Closer analysis of those two scenarios reveals that the initial results provided support for the inclusion of the PoI in all loci but one (D1S1656 in C02 and D3S1358 in H06). In those loci, the calculated LRs were 0. A biased analysis excluding those loci would provide strong evidence for including the PoI (sub-source^2^ log_10_ LR of 27.02 and 23.41, respectively). A skilled user should notice this imbalance between the estimated results for different loci and act accordingly, but the question remains why this happens. Algorithms that consider the genotype set as a parameter of the posterior (like STRmix™) might struggle if the genotype of the PoI is only included in genotype sets that are unlikely under the considered model. It is then possible that no Markov chain will ever explore those genotype sets when an insufficient number of iterations is performed. This does not seem to be specific to the present benchmark, since an independent example of a mixture leading to this problem has been described by Lin et al. [15]. Buckleton et al. [3] noted:

> “We would ask the question: ‘Does one want an *LR* system to allow 15,18 [PoI genotype – our note] in this case?’ If you answer ‘yes’ then the inevitable result is that a much higher false inclusion rate will occur.”

We here show that this is actually avoidable, since the HMC method does not display a higher false inclusion rate than STRmix™ (Table 1), yet provides strong support for the true hypothesis in those two scenarios. Indeed, HMC provides strong inclusion for both scenarios with log_10_ LR of 24.94 for scenario C02 and 20.07 for H06. The problematic loci, however, correctly resulted in low sub-sub-source LRs: 0.0044 for D1S1656 in C02 and 0.000041 for D3S1358 in H06.

This suggests that the behaviour of STRmix™ is caused by a combination of the modelling decision to treat the genotype set as a modelled parameter and running an insufficient number of MCMC iterations. Those decisions were made due to the computational complexity of the inference and can be avoided by, for example, using GPU acceleration [1].

An interesting question is what happens when the PoI’s genotype is represented in the posterior estimate of one of the chains, but absent in the others. In this case, the intra-chain estimate for LR is 0 for all chains except one, but the final answer to the user will still indicate a positive LR. This is an example of the nuances one has to deal with if convergence of the estimated distribution over the discrete genotype sets is not rigorously checked.

### 3.2 Results on the verbal scale

Some sources recommend reporting results on a verbal scale that reflects the strength of the evidence [16, 17, 18]. This removes small, likely technical, differences between analysis methods and is supposably easier to understand in court. However, it also introduces arbitrary thresholds by binning the possible values of LR. We analyse the effect this binning has on the results from the tested analysis methods on two verbal scales: the official SWGDAM scale [18] and an alternative scale suggested by ENFSI [17].

Both scales are compared in Table 2.

**Table 2.** A comparison between the SWGDAM verbal scale and the ENFSI recommendation for reporting LRs. The provided log_10_LR ranges present the scenario of reporting the hypothesis of the prosecutor. As both of the scales do not define precisely how to deal with LRs between below 1, we set an LR of 1.5 as the border between grades A, B, and C.

| SWGDM scale | Grade | $\log_{10}\text{LR}$ | Scale suggested by ENFSI |
| --- | --- | --- | --- |
| Supports alternative | A | $< -\log_{10}(1.5)$ | Supports alternative |
| Uninformative | B | $[-\log_{10}(1.5), \log_{10}(1.5))$ | Does not support any proposition |
| Limited support | C | $[\log_{10}(1.5), 1)$ | Weak support |
| | D | $[1, 2)$ | Moderate support |
| Moderate support | E | $[2, 3)$ | Moderately strong support |
| | F | $[3, 4)$ | Strong support |
| Strong support | G | $[4, 6)$ | Very strong support |
| Very strong support | H | $\geq 6$ | Extremely strong support |

We present 10 contingency tables to compare the results on the verbal scale between HMC and the other softwares (STRmix™ or Euroformix). For each number of contributors, we present two tables: one where *H*_*p*_ with the true contributors is tested, and one where the support for the defendant (i.e. support for *H*_*d*_) is tested for false contributors. As HMC and STRmix™ are based on similar models, the verbal scale classification of their results is similar. On the higher-resolution scale suggested by ENFSI for true hypotheses over 2-contributor mixtures, these two methods produce the same verbal classification in 96.43% of scenarios (297/308, Table 3). For true hypotheses over 3-contributor mixtures, they agree in 92.74% of scenarios (409/441, Table 6) and for 4-contributor mixtures in 88.19% of scenarios (448/508, Table 11). Thus, consensus becomes rarer with larger increasing number of contributors. It is also rarer when the false hypotheses are considered: 93.18% for 2-contributor scenarios (287/308, Table 5), 82.09% (362/441, Table 7) for 3-contributor scenarios, and 70.67% (359/508, Table 12) for 4-contributor scenarios. This is likely because both methods become less precise when LRs are low (cf. Fig. 7) and the genotype set of the suspect is unlikely in at least some of the loci.

**Table 3.**
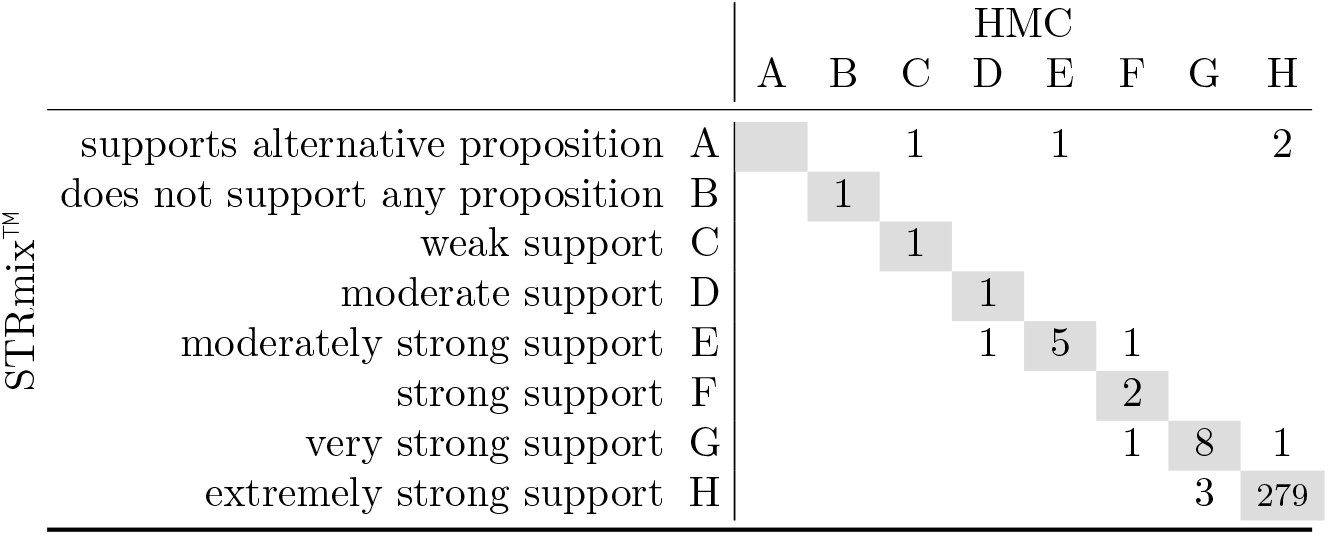
Comparison of the results of HMC and STRmix™ when a verbal scale is used. True hypotheses, 2-contributor scenarios.

**Table 4.**
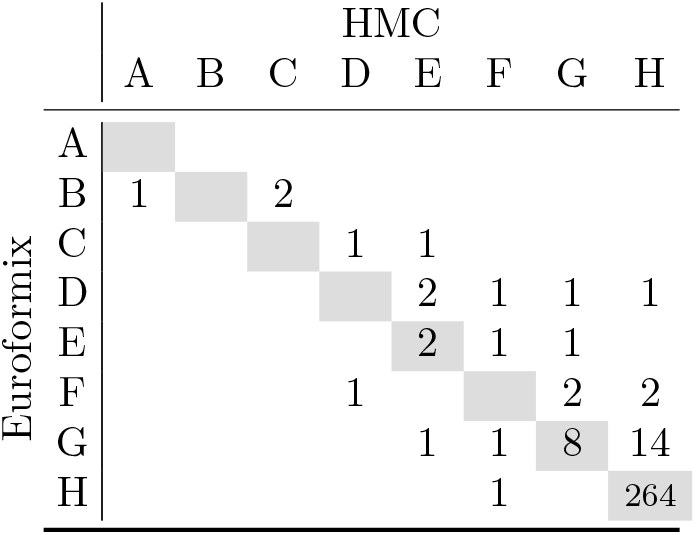
Comparison of the results of HMC and Euroformix when a verbal scale is used. True hypotheses, 2-contributor scenarios.

**Table 5.** Comparison of results of HMC and STRmix™ when a verbal scale is used. False hypotheses, 2-contributor scenarios.

| STRmix<sup>TM</sup> | HMC |  |  |  |  |  |  |  |
| --- | --- | --- | --- | --- | --- | --- | --- | --- |
|  | A | B | C | D | E | F | G | H |
|  | A |  |  |  |  |  |  |  |
|  | B | 1 |  |  |  |  |  |  |
|  | C |  | 1 |  |  |  |  |  |
|  | D |  |  | 4 | 2 |  |  |  |
|  | E |  | 1 |  | 1 |  |  |  |
|  | F |  |  |  | 2 |  |  |  |
|  | G |  |  | 1 | 2 | 2 | 4 |  |
|  | H |  |  | 1 |  | 5 | 5 | 276 |

**Table 6.**
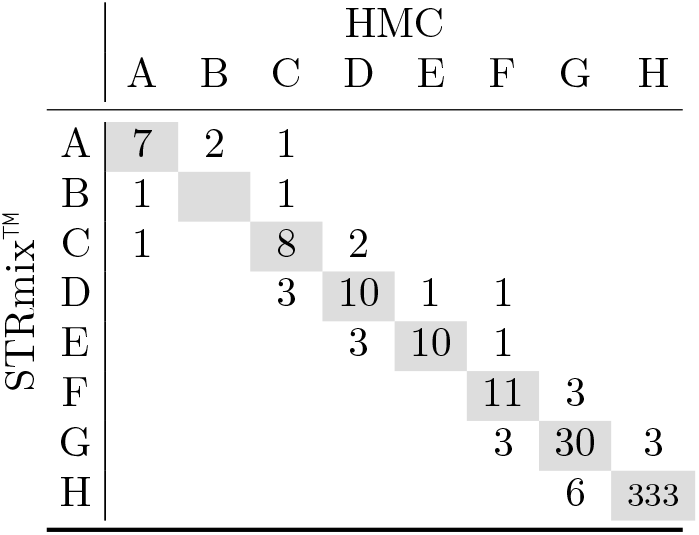
Comparison of results of HMC and STRmix™ when a verbal scale is used. True hypotheses, 3-contributor scenarios.

**Table 7.**
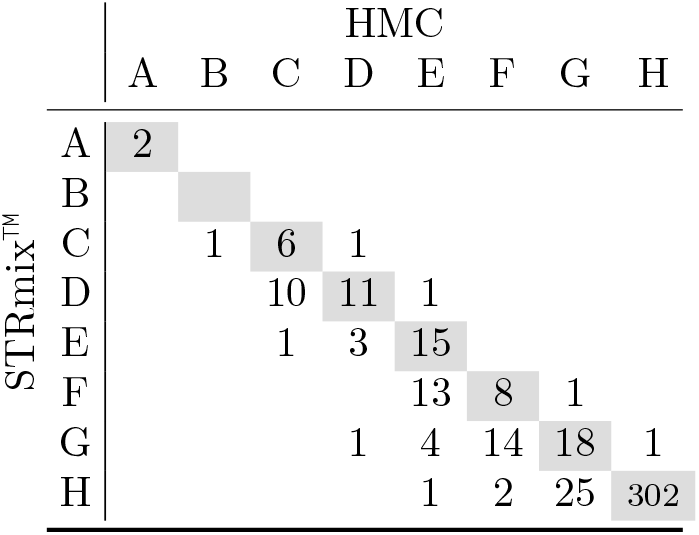
Comparison of results of HMC and STRmix™ when a verbal scale is used. False hypotheses, 3-contributor scenarios.

**Table 8.**
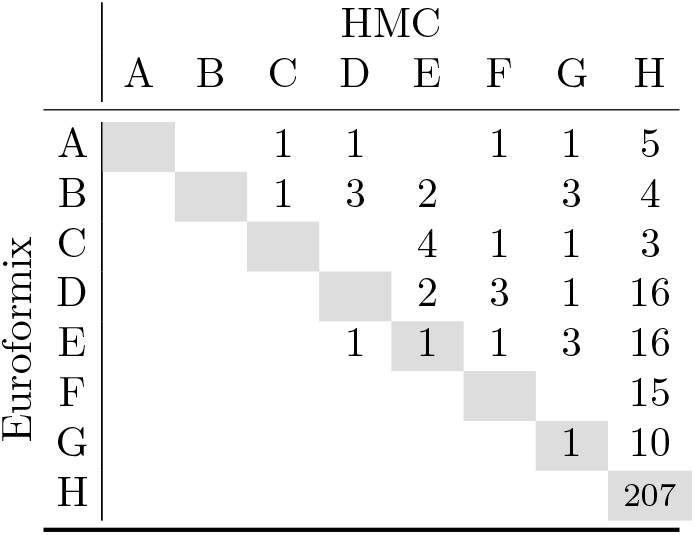
Comparison of results of HMC and Euroformix when a verbal scale is used. False hypotheses, 2-contributor scenarios.

**Table 9.**
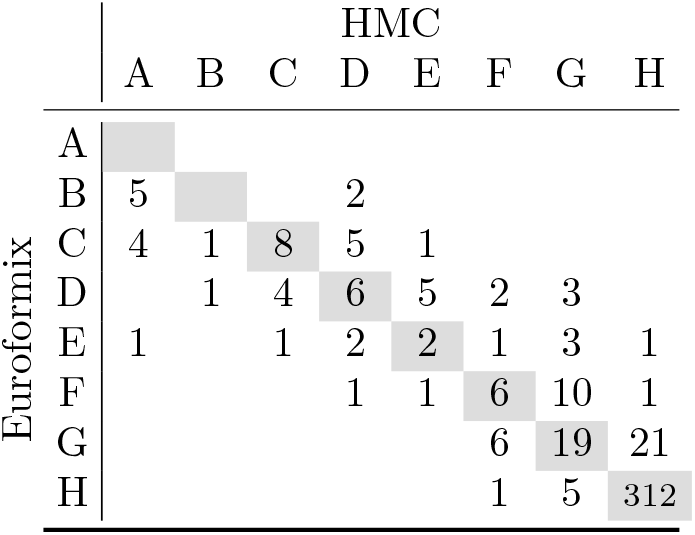
Comparison of results of HMC and Euroformix when a verbal scale is used. True hypotheses, 3-contributor scenarios.

**Table 10.**
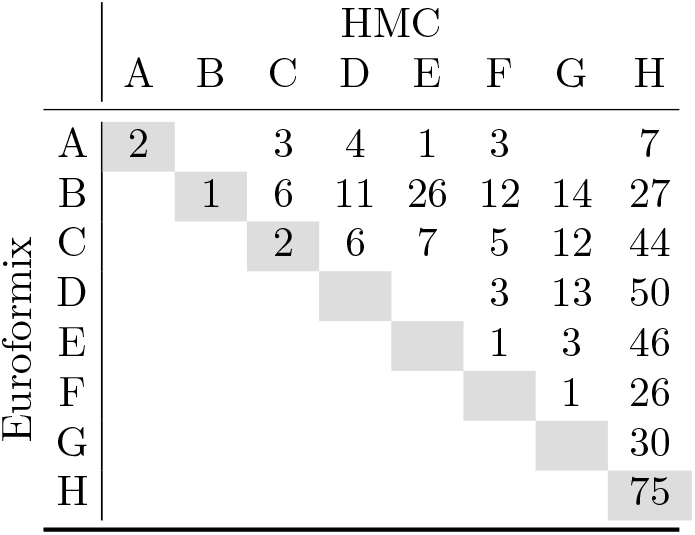
Comparison of results of HMC and Euroformix when a verbal scale is used. False hypotheses, 3-contributor scenarios.

**Table 11.**
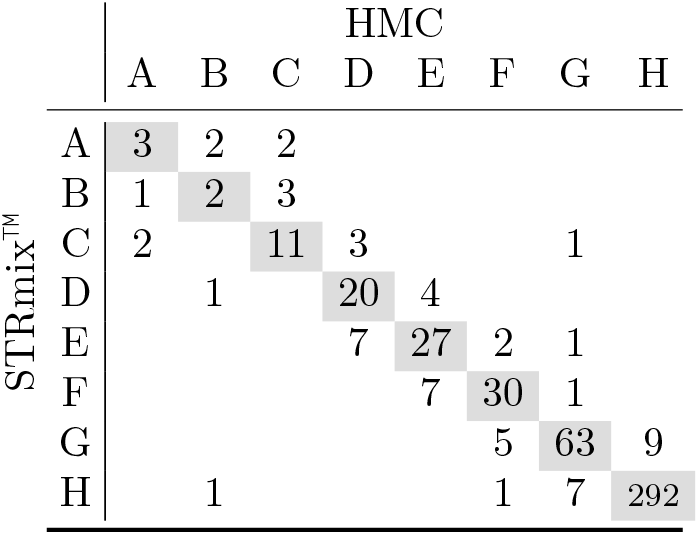
Comparison of results of HMC and STRmix™ when a verbal scale is used. True hypotheses, 4-contributor scenarios.

**Table 12.**
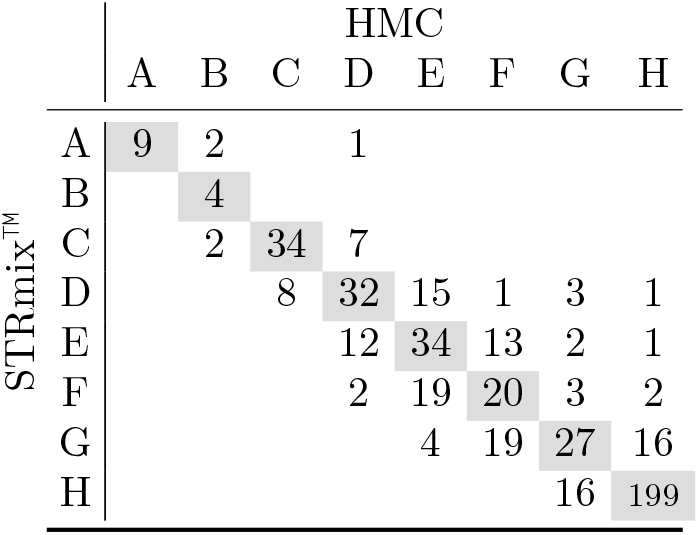
Comparison of results of HMC and STRmix™ when a verbal scale is used. False hypotheses, 4-contributor scenarios.

The discrepancy between HMC and Euroformix is larger, which may be explained by the different probabilistic models, ATs, and optimization formulations (Euroformix has been run in MLE mode, though an MCMC implementation is available, in order to be able to directly compare the results with Riman et al. [2] who also used the MLE mode). In cases where there is a difference in the reported grades, HMC tends to provide stronger support in scenarios with true *H*_*p*_. This is the case for 9.42% (29/308, Table 4) of all 2-contributor scenarios and 12.47% (55/441, Table 9) of 3-contributor scenarios. In the 3-contributor cases, Euroformix provides a stronger verbal equivalent in 33 (7.48%) of cases.

The largest discrepancies can be observed between Euroformix and HMC under the false hypotheses. In many of these cases, HMC provides stronger negative evidence. This is the case in 31.82% (98/308, Table 8) of 2-contributor scenarios and 81.86% (361/441, Table 10) of 3-contributor scenarios. When compared with the original benchmark [2], HMC provides stronger negative evidence in 94.29% (479/508) of the 4-contributor scenarios. The opposite situation, where Euroformix provides stronger negative verbal equivalent, is rare, with only one scenario among 2-contributor ones and no scenarios among 3-contributors ones.

### 3.3 Results on the numerical scale

In order to provide a refined comparison, we plot in Figure 3 all of the computed LRs when true hypotheses were considered. This again highlights that HMC and STRmix™ provide similar results. In 95.54% (1201/1257) of all scenarios the difference between the LRs computed by these two methods is smaller than 2 bans. In 98.97% (1244/1257) of cases the difference is smaller than 4 bans, and only in 3 scenarios the difference is larger than 6 bans. These extreme cases are CO2 and H06, which were already discussed in Ref. [2] and in Subsection 3.1, as well as E05 RD14−0003−33_34_35_36−1;4;4;1−M2a−0.63GF (E05). In scenario E05 with Contributor 33 the PoI, HMC computes log_10_ LR = *−*0.0852, while STRmix™ outputs 6.5106. We analyse this case in detail in Supplementary Material 2.

**Figure 3.**
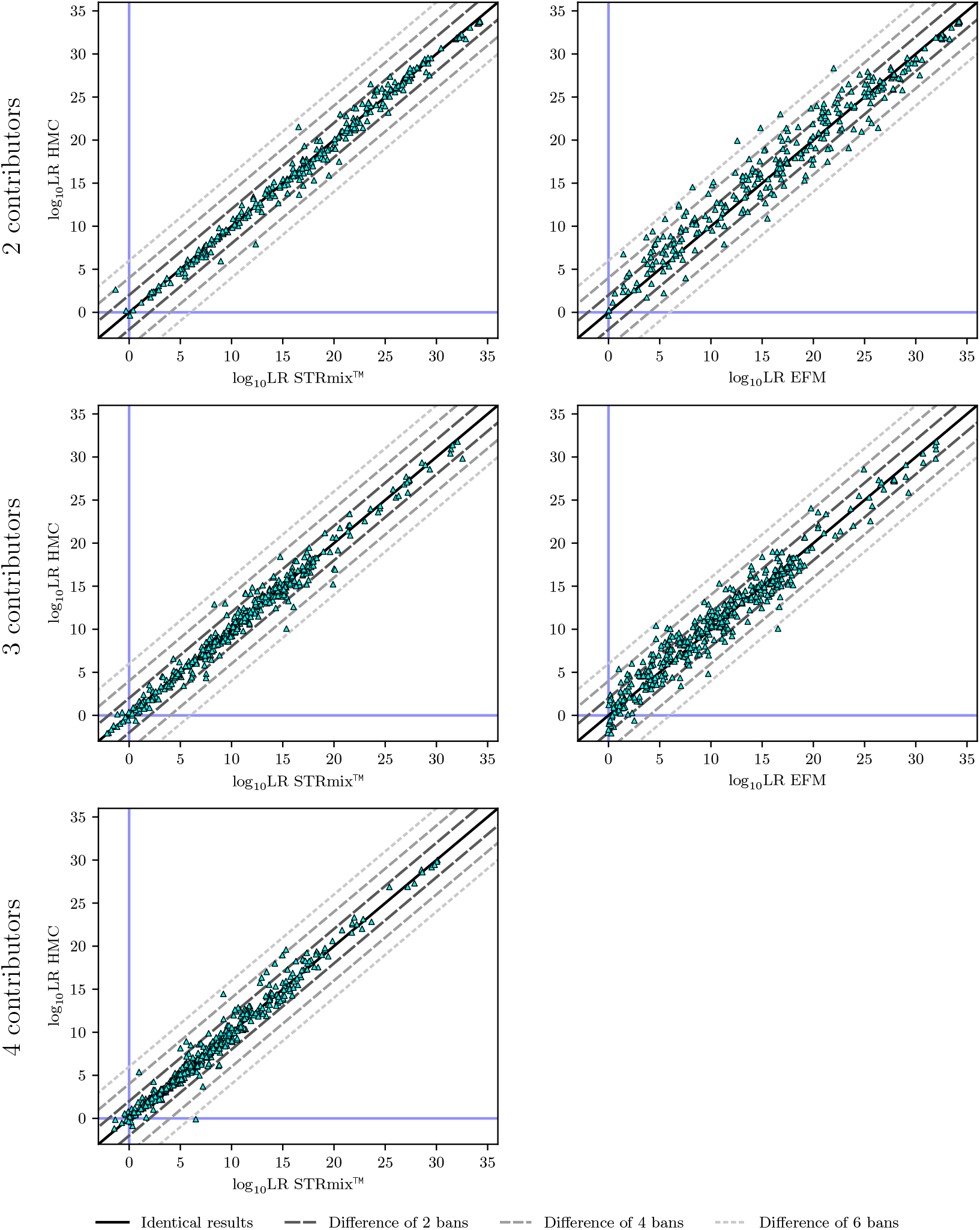
Comparison of the computed LRs for all tested true hypotheses between HMC (*y* axes) and the other tested methods (*x* axes, STRmix™ and Euroformix (EFM)). The two 2-contributor scenarios for which STRmix™ outputs an LR of 0 are omitted from the plots, since log_10_ LR is undefined in those cases.

Larger discrepancies are observed between HMC and Euroformix, with 22.43% (168/749) of scenarios with a true contributor in *H*_*p*_ resulting in LRs that differ by more than 2 bans, 4.01% (30/749) by more than four bans, and 0.8% (6/749) by more than six bans. These differences are visualised again as log_10_ LR histograms in Figure 4.

**Figure 4.**
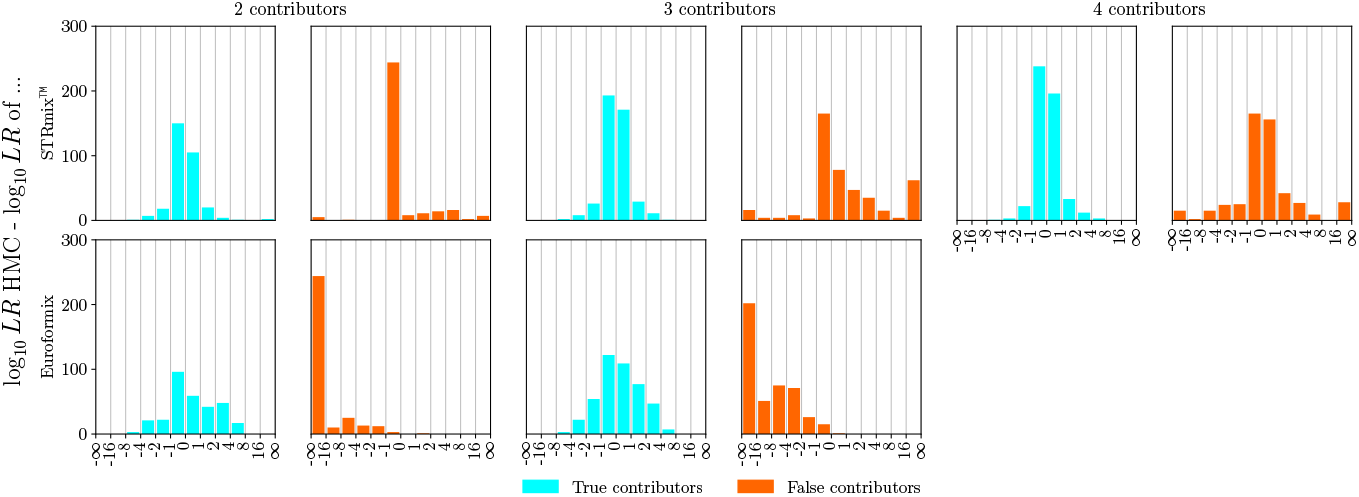
Histograms of the differences in log_10_ LR between HMC, STRmix™, and Euroformix for both true (cyan) and false (orange) hypotheses.

For the false hypotheses, a slight trend toward stronger negative evidence can be observed in the results of STRmix™ (Figure 5). There are 61 2-contributor scenarios where log_10_ LR is lower when STRmix™ is used (and 11 scenarios with the opposite being true), 244 3-contributor scenarios (57 otherwise), and 262 4-contributor scenarios (203 otherwise). We note, however, that the number of scenarios where STRmix™ resorted to trivial rejection of the hypothesis (i.e. LR=0) is larger than for HMC: 202 3-contributor scenarios for STRmix™ vs. 155 for HMC; 71 4-contributor scenarios for STRmix™ vs. 54 for HMC. Two algorithmic factors influence the occurrence of inference results with LR=0. First, the way the considered genotype sets are pruned by removing unlikely ones may lead the algorithm to ignore the genotype of the PoI. The second factor is the general behaviour of MCMC methods to consider a single genotype set at a time, as discussed in Subsection 3.1.

**Figure 5.**
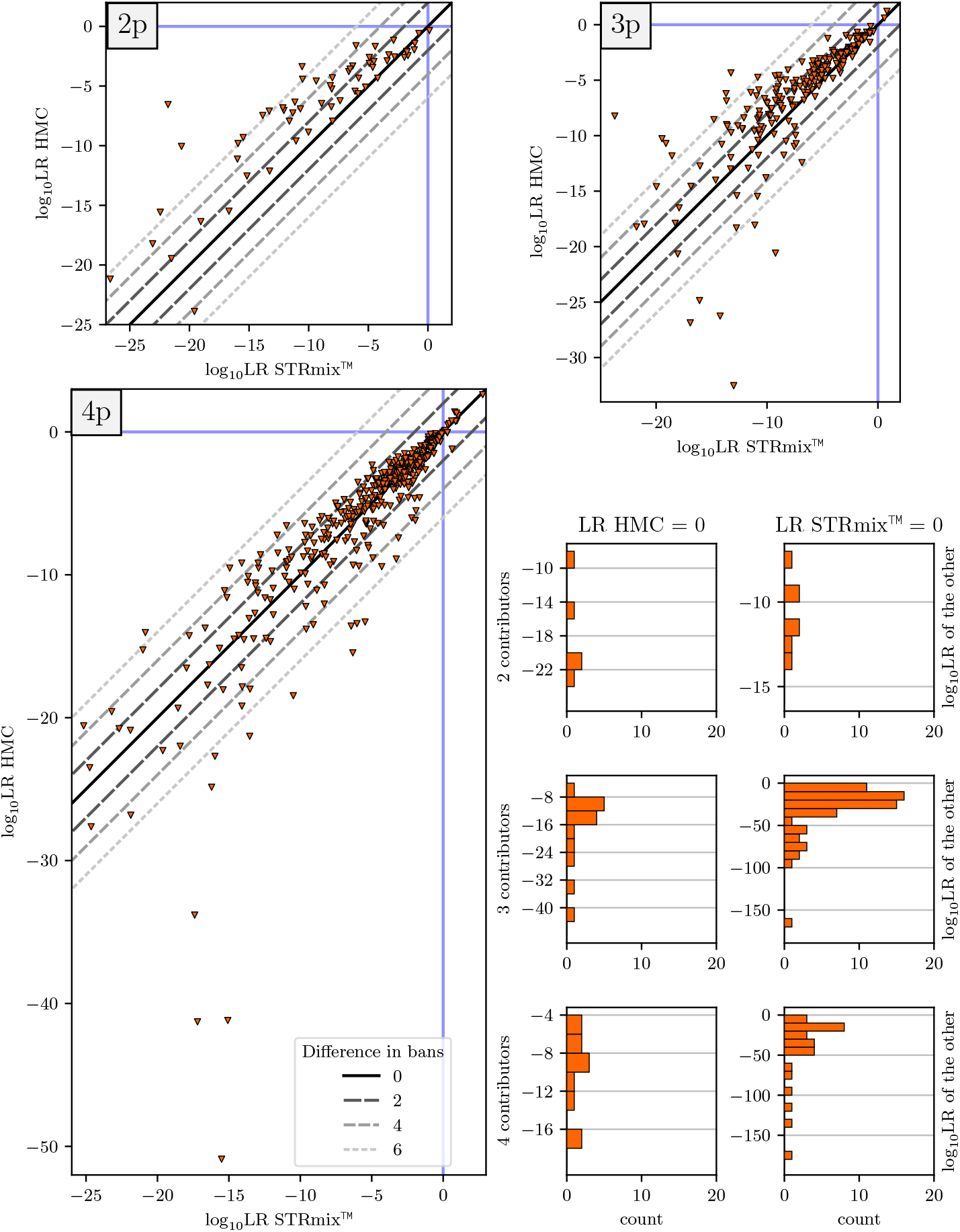
Comparison of HMC and STRmix™ for all false hypotheses. The plots marked with 2p, 3p, and 4p present scenarios with two, three, and four contributors, respectively, for which both methods output non-zero LR. The histograms at the bottom-right show the distributions of LRs from the respective other method in scenarios for which exactly one of the methods outputs LR=0. In addition, there are 236 2-contributor, 140 3-contributor, and 43 4-contributor scenarios were both methods output LR=0.

For scenarios resulting in stronger negative LRs, a larger discrepancy between HMC and STRmix™ is observed. We believe the main reason for this is that the unlikely genotypes of the false contributors are better explained by the tails of the estimated posterior, especially the right tails of the peak-height standard deviation parameters. Any difference between the methods becomes more apparent in the tails and, combined with the lower precision in these cases (see Figure 7), results in more noise.

When comparing HMC with Euroformix, we observe that Euroformix tends to not provide strong negative evidence (Figure 6). The largest differences (if HMC did not result in LR=0) are observed for the scenarios:

**Figure 6.**
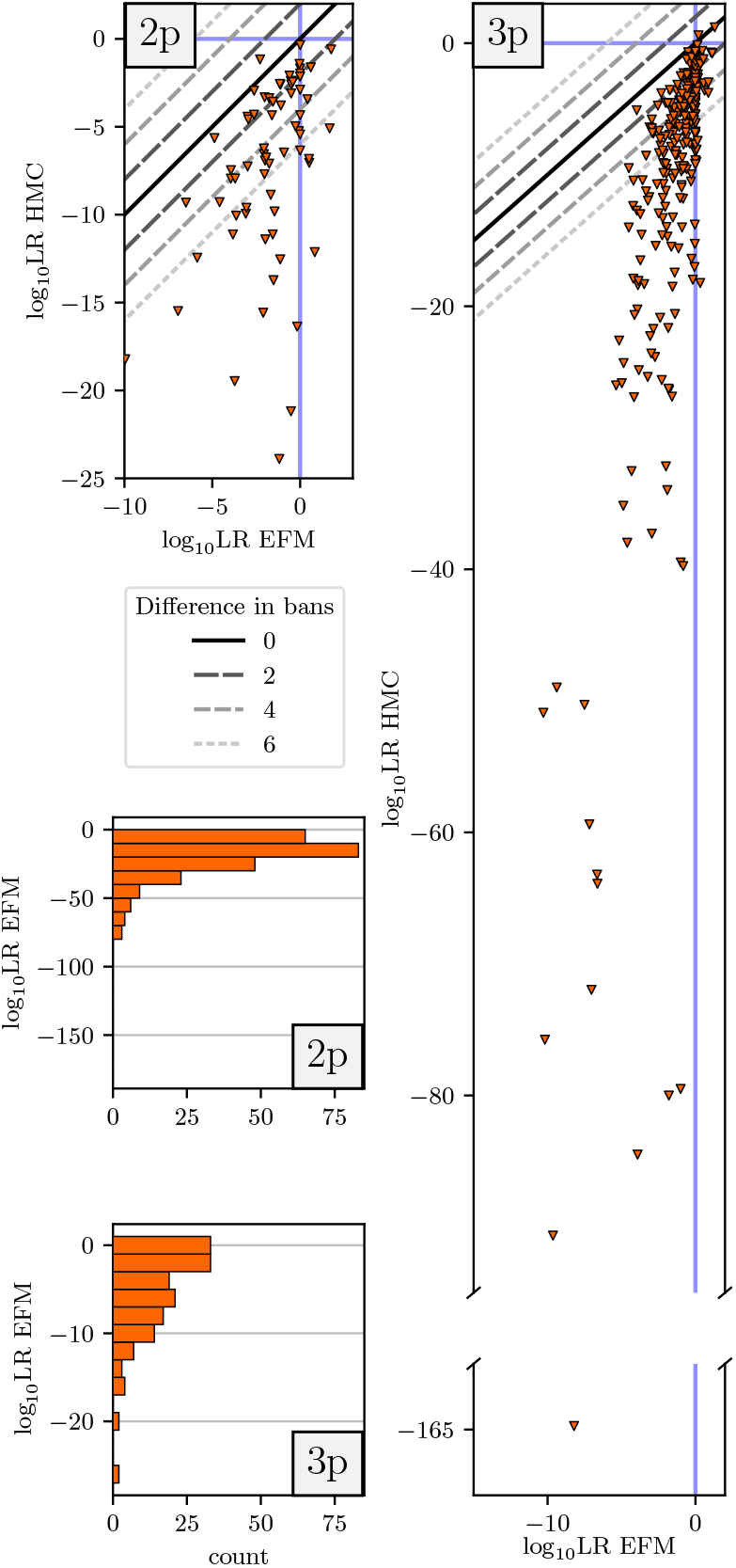
Comparison of HMC and Euroformix (EFM) for all false hypotheses. The plots marked with 2p and 3p present scenarios with two and three contributors, respectively, for which HMC outputs non-zero LR. The histograms at the bottom-left show the distributions of LRs from Euroformix in scenarios for which HMC outputs LR=0. Euroformix outputs non-zero LR for all scenarios in the benchmark.

- 2-contributor: mixture B04 RD14−0003−42 43−1;9−M2U60−0.15GF, PoI is C06C Cauc - Euroformix log_10_ LR = *−*1.19, HMC *−*23.9.
- 3-contributor: mixture B01_RD14−0003−36_37_38−1;2;1−M2a−0.5GF, PoI is PT84183_AA - Euroformix log_10_ LR = *−*8.21, HMC *−*164.73.

However, this would be important only in cases in which the expert has to report the strength of the negative evidence.

### 3.4 A new precision benchmark

Most previous studies [2, 5] compared methods by running them once on every scenario. Two notable exceptions are the study by Bright et al. [19], in which STRmix™ has been tested multiple times on a set of in-house 2-contributor mixtures, and the study by Riman et al. [2], where the computations for 8 scenarios were repeated twice. Among these 8 scenarios were the two scenarios described in Subsecion 3.1 and 6 scenarios in which the convergence metric of STRmix™ remained above the threshold of 1.2 suggested by the authors of the algorithm. In those 6 scenarios the average difference between the two runs was 0.282 bans [2]. The inverse of the run-to-run variability over multiple runs with identical hyper-parametrisation on the same EPG data for the same hypotheses defines the *precision* of a probabilistic genotyping method. Precision has to be considered in addition to accuracy [20].

In order to quantify the precision of the HMC method, we here introduce a new benchmark that expands upon earlier results [1]. Because precision studies of other methods are not available, we cannot provide a comparison. In order to keep the benchmark tractable, we select 23 mixtures uniformly at random (7 2-contributor, 6 3-contributor, and 10 4-contributor mixtures)^3^. For each mixture, we run HMC inference for all scenarios provided in Ref. [2], thus a total of 144 scenarios. We run each scenario 10 times, resulting in 1440 runs, and display the resulting LRs in Figure 7. In one of the scenarios, HMC provides an OotNT result: the 4-contributor scenario G06 results in log_10_ LR between *−* 0.937 and *−* 0.637 when Contributor 33 is the PoI.

**Figure 7.**
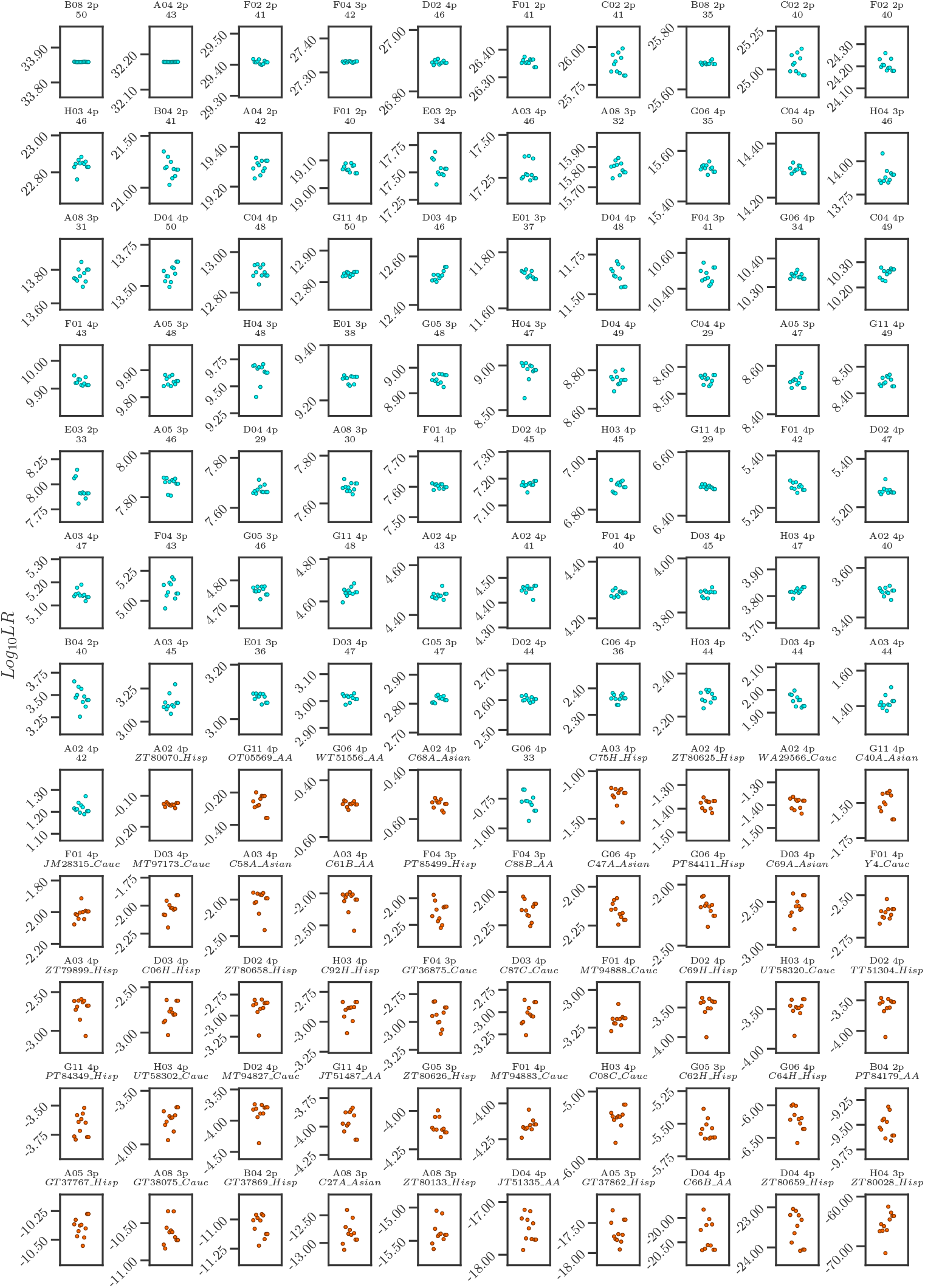
Run-to-run variability of HMC. Each small plot shows the log_10_ LR computed across 11 runs for one of the 120 scenarios (see Supplementary Material 1 for the list of scenarios). True hypotheses are plotted in cyan, false ones in orange. 24 scenarios that resulted in LR=0 in all 10 runs are skipped.

We quantify the precision of HMC inference by the standard deviation of the log_10_ LR and their min–max span (Figure 8). Supporting the hypothesis set out in the Supplementary Material of Ref. [1], we generally observe lower precision when considering false hypotheses. The average ± standard deviation of the standard deviations of the log_10_ LR across all 2-contributor scenarios was 0.0415 ± 0.0372, 0.0335 ± 0.0293 for 3-contributor scenarios, and 0.0213 ± 0.0156 for 4-contributor scenarios. The outlier among 3-contributor scenarios with false hypotheses is the case with the lowest estimated log_10_ LR (excluding cases for which all 11 runs result in LR=0): H04 3p with ZT80028 Hisp as the PoI. In this case, all per-run log_10_ LR *<−*60. Taken together, these results confirm the high precision of the HMC method [1]. In all considered cases, the min–max span of the LRs in scenarios with true hypotheses was smaller than 0.5 bans with standard deviation below 0.1 bans.

**Figure 8.**
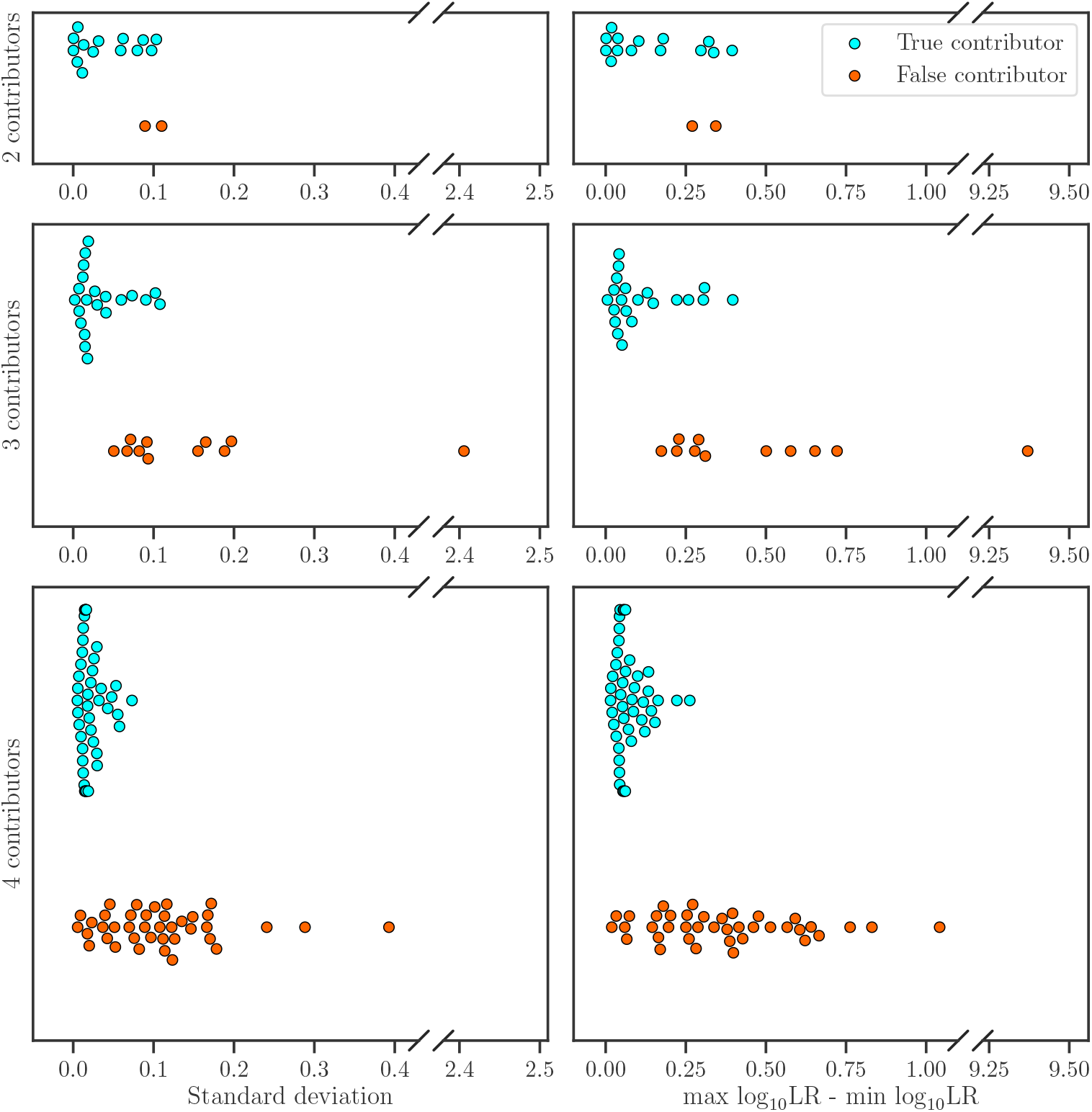
Standard deviations and min–max spans of the log_10_ LR values in all scenarios of the precision benchmark across 11 repeated runs of the HMC method for each scenario. The presented results also include Samples 1b and 2b from Ref. [1]. Scenarios with LR=0 are omitted.

Next, we count how often the result on the ENFSI verbal scale (cf. Subsection 3.2) changed between repeated runs on the same scenario. Considering the grades assigned to the prosecutor’s hypotheses, 48 scenarios are classified as *extremely strong support*, 9 scenarios as *very strong support*, and 6 as *strong support*, 4 as *moderately strong support*, and 3 as *moderate support* unanimously in all 11 runs. The only case where the grades differed between runs of the HMC method was mixture D03 4p with Contributor 47 as PoI: 9 runs classified it as *strong support*, but 2 runs as *moderately strong support*. In this case, however, avoiding the variance between the verbal strength categories is hard, given that HMC estimates log_10_ LR very close to the scale threshold of 3: the lowest LR is 2.9864 and the highest 3.0287, the difference between them is only 0.0423 of a ban. None of the benchmark scenarios was classified as *weak support*. One scenario (A02 4p with false contributor ZT80070_Hisp) unanimously resulted in LRs in all 11 runs that do not support any hypothesis. All other scenarios, including the false negative in one scenario from the G06 4p mixture, unanimously provided support for the alternative hypothesis in all 11 runs. We conclude that a situation in which HMC provides different verbal equivalents when run multiple times is rare, with only one (out of 120) case observed in our benchmark.

Finally, we also measure the computer time spent by the HMC algorithm on each of the considered scenarios (Figure 9). Every 3-contributor case was solved in under 11 minutes, and every 4-contributor case in under 1 hour and 3 minutes. These times include the compilation of the coxomputational graph by the Accelerated Linear Algebra library [21] used in HMC.

**Figure 9.**
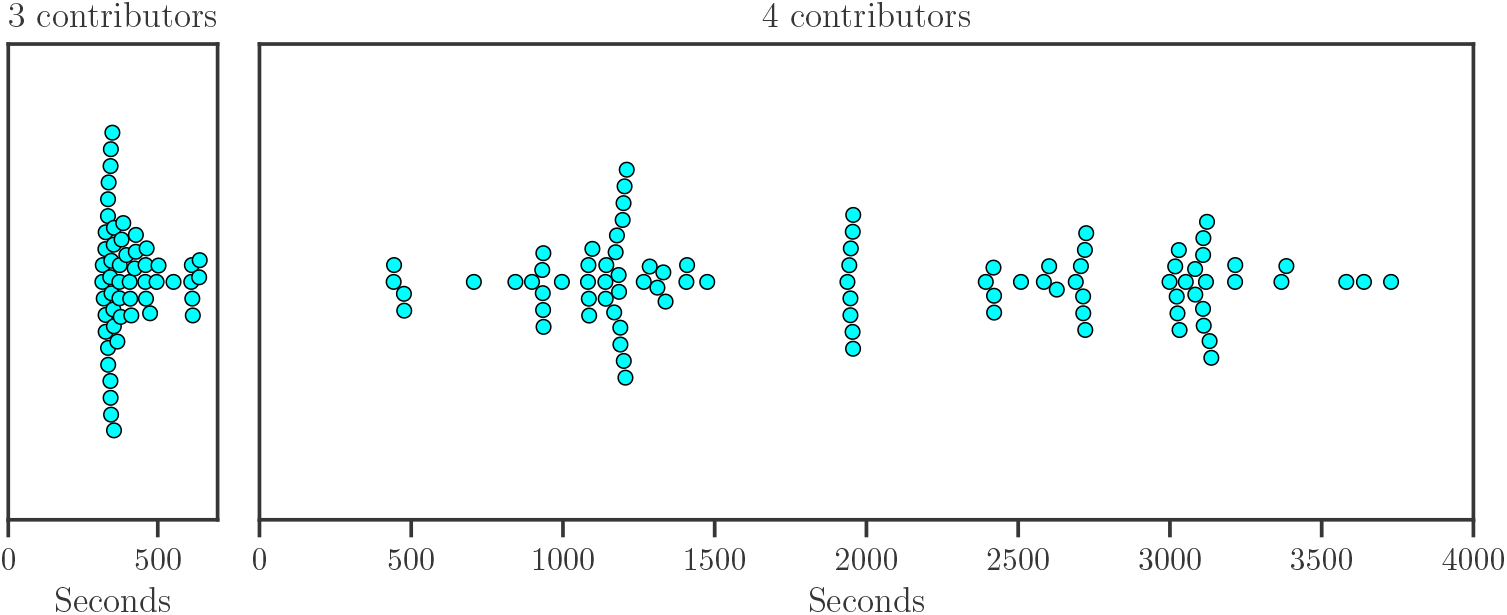
Execution times of the HMC method. Each dot represents a single run of one of the scenarios from the present benchmark. All runs were performed on a single Nvidia Tesla T4 GPU.

### 3.5 Differences between rare-allele models

Buckleton et al. [3] suggested that different rare allele treatment might distort the results of method comparisons (*“There was no compensation for the different rare allele models used by each software*.*”* [3]). We therefore quantify the effect of the rare-allele model by recalculating all LRs of HMC with the posterior mean allele frequency model by Triggs and Curran [22], reusing the deconvolution provided by the previous HMC inference. We then compare the results with LRs obtained using the 5/2N rare-allele model [23]. Both sets of results are calculated assuming contributors from the NIST-1036 Caucasian population [24]; therefore, N=361. We provide a list of the rare alleles within the true contributors according to the NIST-1036 Caucasian population in Supplementary Material 1, tab “Rare contributor alleles”. Using the example of the STRidER’s [25] European population, we also show that the number of rare alleles drops when a larger population sample is used.

The results in Figure 10 show that different rare-allele models lead to differences in log_10_ LR that are larger that the run-to-run variability and therefore should be considered significant. The data points for true contributors can be visually clustered into three categories: PoIs with no rare alleles present in their genotypes, PoIs with one rare allele, and PoIs with two rare alleles. The majority of points from these clusters lie around the horizontal line at 0 for cases with no rare alleles, 0.22 for cases with one rare allele, and 0.44 for cases with two rare alleles. To be precise, the 25th and 75th percentiles of the differences on the logarithmic scale are 0.221… 0.439 for true contributors with two rare alleles, 0.013 … 0.226 for true contributors with one rare allele, and 0.000 … 0.017 for true contributors with no rare alleles. The source of this behaviour might be sought in the estimated rare allele frequencies. Let us consider the value of 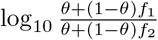, where *f*_1_ is the rare allele frequency from the 5*/*2*N* model, from the posterior mean mean allele frequency model and *θ* is the co-ancestry correction factor. This value is close to 0.22 for all of the rare allele loci: for SE33 the value is 0.225, for D22S1045 0.223, and for TPOX 0.219 with 39, 15, and 8 distinct allele scores in the population, respectively.

**Figure 10.**
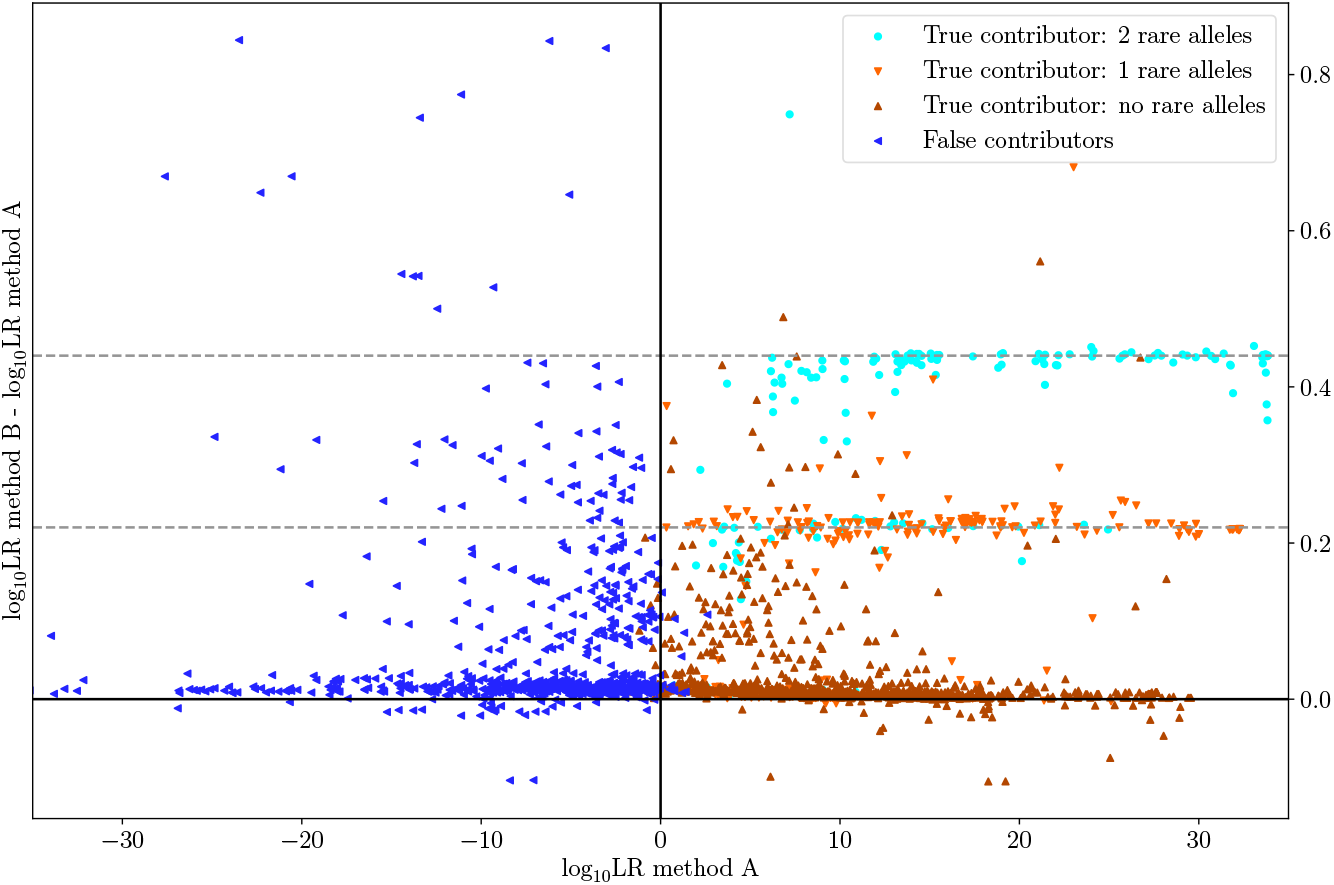
Difference between the log_10_ LR obtained by HMC with two different rare-allele models: the 5/2N model (Method A) and the posterior mean allele frequency model (Method B). Colours denote whether the PoI is a true contributor and, if so, how many rare alleles are present in his/her genotype. The dashed horizontal lines are at differences of 0.22 and 0.44. The true hypotheses with PoIs with rare alleles cluster around those lines.

Points away from the respective line of their cluster are often caused by the rare allele(s) dropping out from the mixture. This was often the case especially for alleles 19.2 and 28 in the FGA locus and for Contributor 50’s rare allele 7 in the TPOX locus. The corresponding markers are located towards the right ends of the dyes, such that these peaks risk to drop out in degraded mixtures.

When there are rare alleles present in both the PoI’s genotype and the mixture, the log_10_ LR difference between the 5/2N model and the posterior allele frequency model will depend on the value of N in the 5/2N formula and therefore on the population allele frequencies. For different values of N, the data points will cluster around horizontal lines at different values. Following the same reasoning as above, the points would be expected to cluster around 0.52 for N=100, and around 0.095 for N=1000.

### 3.6 Differences between two settings of Euroformix

As discussed in Subsection 2.3, our benchmark setting required raising the analytical thresholds. This must be done in order to avoid double-backward stutters that are not modelled by Euroformix. In case any double-backward stutter was present in a profile, it could significantly distort the result. When the analytical threshold (AT) is raised, we expect the results to express lower confidence (i.e. lower LRs for true contributors, higher LRs for false contributors). This is because less information is retained from the mixture at higher ATs, as analysed in Supplementary Material 3. We therefore compare the results obtained by Riman et al. [2] with Euroformix version 2.1.0 at lower ATs with the results we obtained here with Euroformix version 3.4.0 and higher ATs. The comparison is presented in Figure 11. As expected, higher ATs mostly lead to lower strengths of evidence. The exceptions are the 2-contributor mixtures with forward stutter peaks. Since these stutters are not modelled in Euroformix 2.1.0, the resulting deconvolution must be sub-optimal. This behaviour has independently been observed by Cheng et al. [5]. While the use of higher ATs might be perceived as a disadvantage for Euroformix, we note that this was required because of the limitations of the model. An alternative benchmark design would involve all the methods tested with higher ATs.

**Figure 11.**
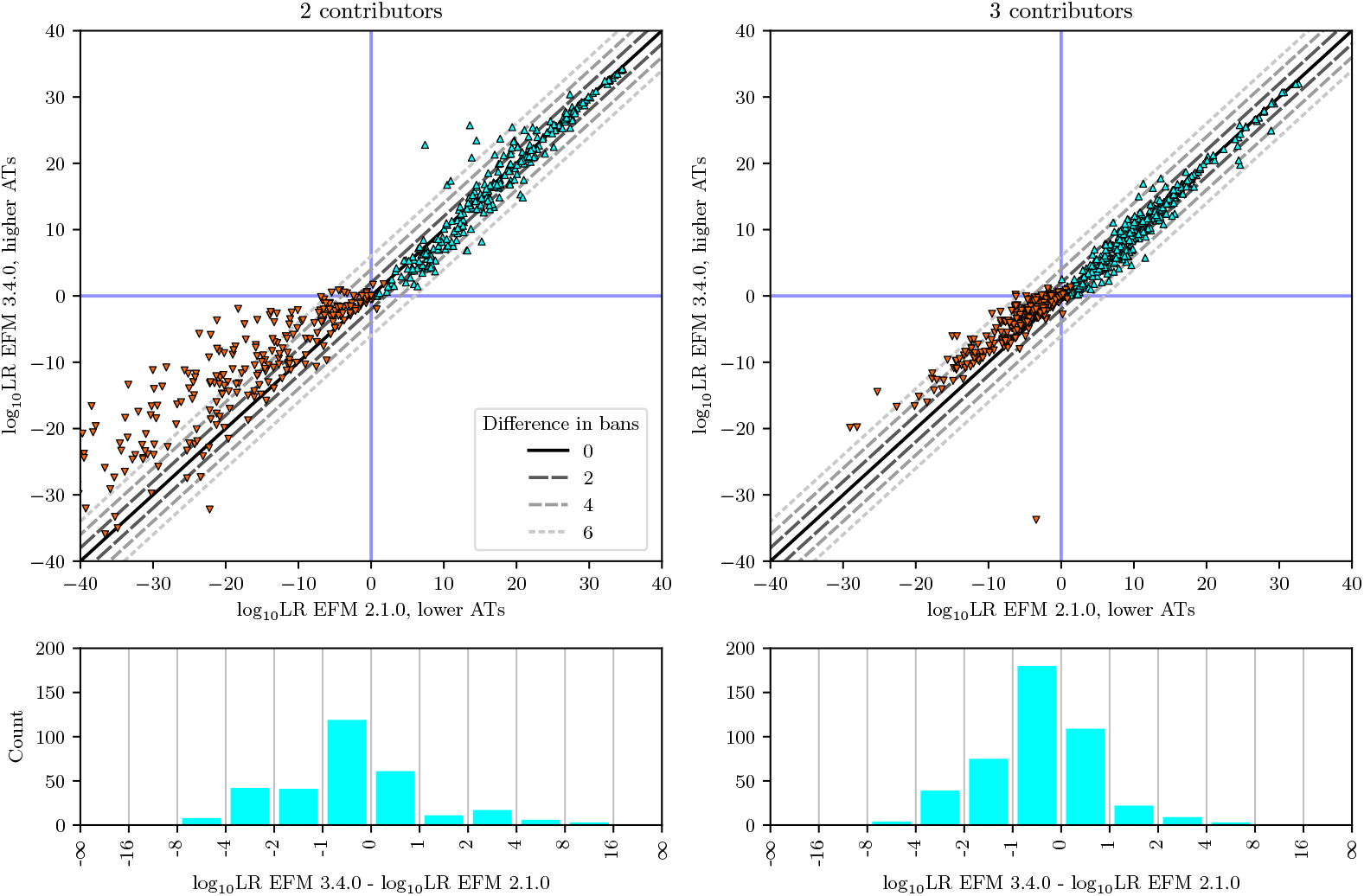
Difference between the results obtained with versions 2.1.0 and 3.4.0 of Euroformix (EFM) at different analytical thresholds (AT). **Top row:** Scatter plots of the scenarios for 2-contributor (left) and 3-contributor (right) mixtures. True hypotheses are in cyan, false ones in orange. Scenarios where any of the software versions computed log_10_ LR below *−*40 are omitted. **Bottom row:** Histograms of the differences for all scenarios with true PoI in 2-contributor (left) and 3-contributor (right) mixtures.

## 4 Conclusions

We provided a large-scale validation study of the HMC method for probabilistic genotyping [1] and compared it with the current state of the art. The results revealed that the HMC method achieves state-of-the-art classification performance at high precision (i.e., low run-to-run variability). In OotNT scenarios, HMC did not provide large positive or negative LR values, and correctly manifested uncertainty.

The classification results obtained by HMC were similar to those obtained using the popular software STRmix™. The similarity can be explained by the fact that the probabilistic genotyping model used by HMC is based on the work by Taylor et al. [11]. Compared to Euroformix, HMC exhibited a lower OotNT rate, a significantly better ability to provide negative evidence, and a slightly higher area under the ROC curve for 3-contributor mixtures.

We further introduced a new precision benchmark that can serve as a baseline for future methods development and comparison. This benchmark provided additional results corroborating the high precision of HMC, which stems from the fact that this algorithm considers genotype sets in every iteration and assures proper chain convergence.

In Supplementary Material 3, we analysed the factors influencing the strength of the evidence provided by the HMC method and how these factors correlate with algorithmic differences between HMC and the other algorithms. We find that the information content of the short tandem repeat mixtures influences the confidence of the HMC algorithm, corroborating earlier results [5, 26]. This provided additional support that the strength of the evidence decreases with:

- decreasing total amount of DNA material in the mixture,
- decreasing amount of DNA material coming from the PoI,
- decreased weight of the contributor, and
- decreased sample quality (i.e., increased “quality index”).

While this study focused on understanding the differences between the algorithms, all of the compared methods are characterised by high accuracy and a healthy ability to provide a lower strength of evidence when the information content in the electropherogram is lower. We note, however, that our approach of analysing all mixtures of the benchmark might not directly reflect individual casework pipelines. This is because:

- Laboratories might impose additional conditions on the quality of a mixture. If these conditions are not satisfied, a mixture might not be analysed at all or not reported, or a different replicate might be used. We discussed an example of such an issue with one of the mixtures in Supplementary Material 2.
- Human expert analysts will scrutinise the results output by an algorithm. For example, in the case of the two scenarios discussed in Subsection 3.1, an experienced analyst would immediatelty notice the unnatural distribution of per-locus LRs.
- Despite great efforts by the PROVEDit team to create a plausible laboratory simulation of sample degradation, there remains the possibility of a covariate shift between the crime-scene samples and the ones created in a laboratory.

An interesting extension of the benchmarks would be to scenarios in which the (simulated) relatives of the true contributors are considered as PoIs. Such experiments would be more challenging for probabilistic genotyping methods. Initial analyses, though without comparing different models, have already been performed [27].

## Acknowledgments

We thank Dr. João Nunes and Dr. Suzanne Walz (both Biotype GmbH, Dresden) for sharing their insights about the influence of the treatment type on the compromised whole-blood mixtures and Dr. Øyvind Bleka (Oslo University Hospital, Oslo) for his advice on proper usage of the Euroformix software and input on the manuscript.

## Footnotes

1 Following Slooten’s argument, depending on the dataset, the maximum practically achievable value is slightly lower than 1.0.

2 Assuming the other, unreported sub-sub-source LRs excluding the problematical single loci were 0.

3 The list of mixtures and the detailed results are available in Supplementary Material 1.

