## Supplementary Material 3 for "Analysis of the Hamiltonian Monte Carlo genotyping algorithm on PROVEDIt mixtures including a novel precision benchmark"

—

—

### Effects of information content of DNA mixtures on LRs

It has been shown that LR values depend on the amount of DNA material and on the number of contributors [3]. We expect that these observations generalise to fully continuous models and therefore explore how these factors, together with other available information about the mixtures, influence the results. Specifically we focus on the influence of

- the total template mass,
- the template mass from the POI,
- the POI's level of contribution to the mixture (major, minor, equal contributor), and
- the ratio of the concentrations of 80 and 214 base-pair fragments, which is an indicator for the inverse DNA extract quality, referred to as "quality index" (QI)

on the computed LRs in cases where the true contributor is the POI. This is possible because the PROVEDIt dataset contains the required metadata about the laboratory methods and the conditions used to create the mixtures. We also analyse how these factors correlate with differences in the LRs computed with the three tested methods. We will use notation "a (b ... c)" to denote the median $a$, the 25th quantile $b$ and the 75th quantile $c$.

Figure 1: Distributions of the $\log_{10}$ LR computed by HMC when exploring true hypotheses using different amounts of DNA material from the suspect (measured in picogram, pg) for different numbers of contributors.

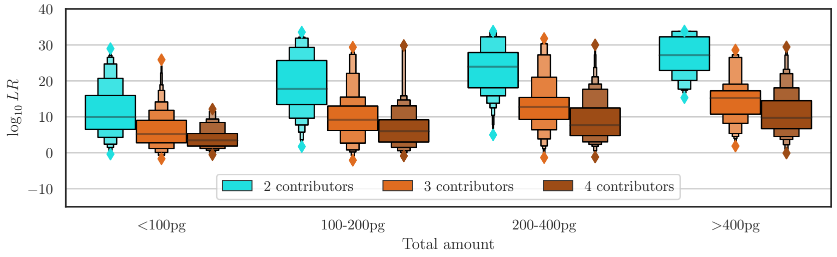

Figure 2: Distributions of the differences between the $\log_{10}$ LR obtained by different methods when true hypotheses are explored with different total amounts of DNA material (measured in picogram, pg) in mixtures with different numbers of contributors.

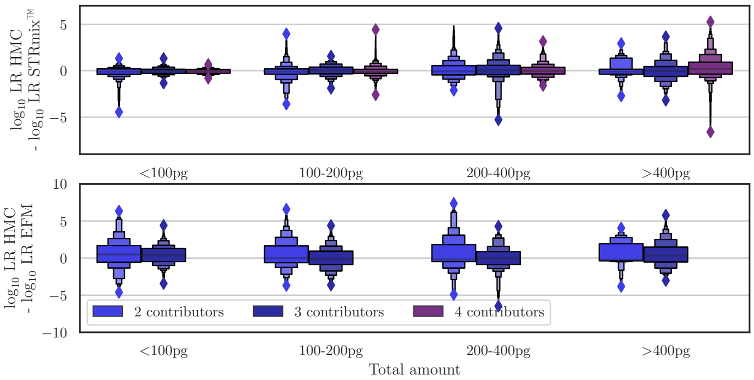


The HMC results are in line with the observation of Taylor [3] that an increasing amount of DNA material increases the LR (Figure 1). This correlation is present regardless of the number of contributors in the mixture. The median $\log_{10}$ LR for 2-contributor low-template (<100 pg) mixtures is 9.92 (6.7 ... 15.94). For high-template mixtures (>400 pg) the median is 27.08 (22.89 ... 32.28). The median for 4-contributor mixtures is 4.14 (1.88 ... 5.35) for low-template scenarios and 11.02 (6.7 ... 14.49) for high-template scenarios.

Next, we study the influence of the total template mass on the differences in LR computed by different methods (Figure 2. Comparing HMC with STRmix™, the median differences are generally small: −0.058 (−0.316 ... 0.166) for low-template mixtures, −0.048 (−0.315 ... 0.277) for 100–200 pg mixtures, 0.002 (−0.404 ... 0.517) for 200–400 pg mixtures, and 0.011 (−0.418 ... 0.73) for high-template mixtures. The largest deviations of the median from 0 are observed in the 4-contributor scenarios. In these cases, STRmix™outputs larger $\log_{10}$ LR for low template mixtures with a median difference of −0.117 (−0.295 ... 0.145), whereas HMC outputs larger $\log_{10}$ LR for high-template mixtures with a median difference of 0.204 (−0.367 ... 0.921).

Comparing HMC with Euroformix (EFM), the median of the differences is 0.417 (−0.5 ... 1.508) for low-template mixtures, −0.059 (−0.809 ... 1.36) for 100–200 pg mixtures, −0.108 (−0.642 ... 1.25) for 200–400 pg mixtures, and 0.18 (−0.437 ... 1.542) for high-template mixtures. The distributions are asymmetric to one side, indicating a prevalence of cases in which HMC outputs larger LRs. This asymmetry become obvious when comparing the medians with the following means: 0.501 for low-template, 0.229 for 100–200 pg, 0.268 for 200–400 pg, and 0.515 for high-template mixtures.

When considering the amount of DNA material from the POI, rather than the total template mass, the correlation with $\log_{10}$ LR is strong, too (Figure 3). In the 2-contributor scenarios, for example, mixtures with low POI template ($\text{template}_{\text{POI}} < 20\,\text{pg}$) result in a median $\log_{10}$ LR of 7.198 (5.533 ... 9.886), while mixtures with high POI template ($\text{template}_{\text{POI}} > 160\,\text{pg}$) result in a median $\log_{10}$ LR of 32.218 (27.634 ... 32.308). The template mass from the POI, $\text{template}_{\text{POI}}$, is calculated as:

$$\text{template}_{\text{POI}} = \frac{\text{total amount} * \text{mixture ratio}_{\text{POI}}}{\sum_i \text{mixture ratio}_i}\,. \tag{1}$$

The differences between HMC and STRmix™show no significant dependence on the amount of DNA from the POI (Figure 4). The only scenarios in which the median deviates from 0 by more than 0.2 are the 4-contributor mixtures with high POI template. There, the median difference in $\log_{10}$ LR is 0.261 (−0.286 ... 0.82). Euroformix provides higher $\log_{10}$ LR than HMC when the template mass from the

Figure 3: Distributions of the $\log_{10}$ LR computed by HMC when exploring true hypotheses with different amounts of DNA material (measured in picogram, pg) from the POI in mixtures with different numbers of contributors.

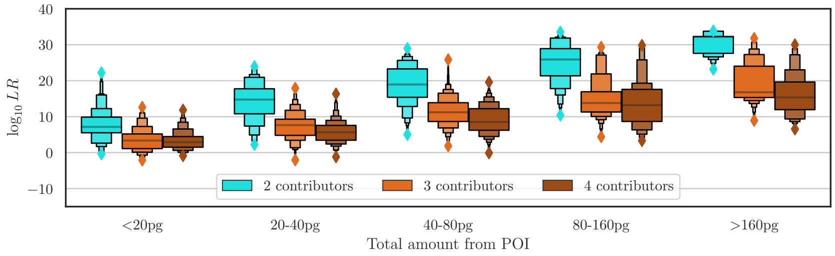


Figure 4: Distributions of the differences between the $\log_{10}$ LR obtained by different methods when true hypotheses are explored with different total amounts of DNA material (measured in picogram, pg) from the POI in mixtures with different numbers of contributors.

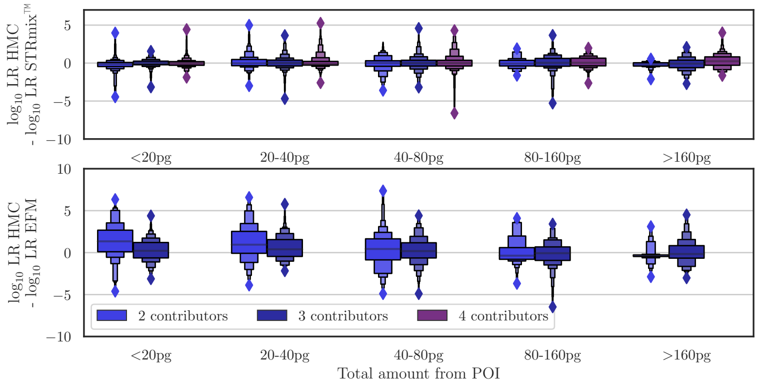

Figure 5: Distributions of the $\log_{10}$ LR computed by HMC when exploring true hypotheses for different types of POIs (minor, equal, major) with different numbers of contributors.

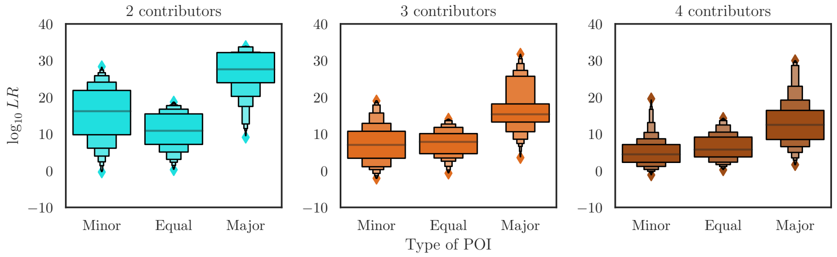


Figure 6: Distributions of the differences between the $\log_{10}$ LR obtained by different methods when true hypotheses are explored for different types of POIs (minor, equal, major) with different numbers of contributors.

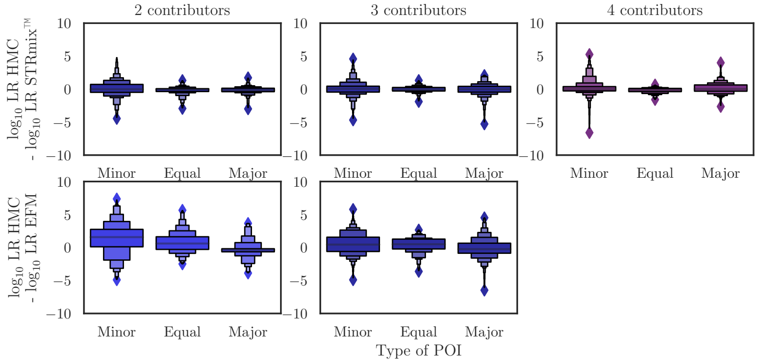


true POI is large. When the total amount of DNA from the POI is 80–160 pg, the median difference is −0.325 (−0.873 … 0.677), and when it is larger than 160 pg the median difference is −0.284 (−0.559 … 0.399). In the other cases, HMC provides stronger evidence: 0.646 (−0.385 … 1.784) in cases with <20 pg DNA from the POI, 0.722 (−0.32 … 1.739) for 20–40 pg, and 0.292 (−0.74 … 1.354) for 40–80 pg.

We repeat the analysis after classifying the benchmark scenarios according to the different contributor types (minor, equal, major) of the POI. Consistently, higher $\log_{10}$ LR are observed when the POI is a major contributor (Figure 5). We compare the LRs obtained when the hypothesis included the true major contributor with the other scenarios. For 2-contributor mixtures, the scenarios with a major POI contribution result in a median $\log_{10}$ LR of 27.638 (24.056 … 32.218), while the median of the minor and equal contributions pooled is 13.843 (8.000 … 17.957). For 3-contributor mixtures, the median is 15.376 (13.257 … 18.2) when the POI is a major contributor and 7.432 (4.083 … 10.445) otherwise. For 4-contributor mixtures, it is 12.496 (8.514 … 13.049) for major POI contributions and 4.897 (2.602 … 7.632) otherwise.

While the differences between HMC and STRmix™do not show any significant trend, Euroformix provides higher $\log_{10}$ LR more often when the POI is a major contributor to the mixture and lower in the other cases (Figure 6). The medians are: −0.326 (−0.82 … 0.451) for major POI contributions, 0.495 (−0.265 … 1.373) for equal contributors, and 0.782 (−0.433 … 2.058) in case of minor POI contributions. The means of the differences are −0.18, 0.564, and 0.829, respectively.

In summary, the results for varying total DNA template mass, POI template mass, and the POI's

Figure 7: Distributions of the $\log_{10}$ LR computed by HMC when exploring true hypotheses for mixtures with different PROVEDIt quality indices and different numbers of contributors.

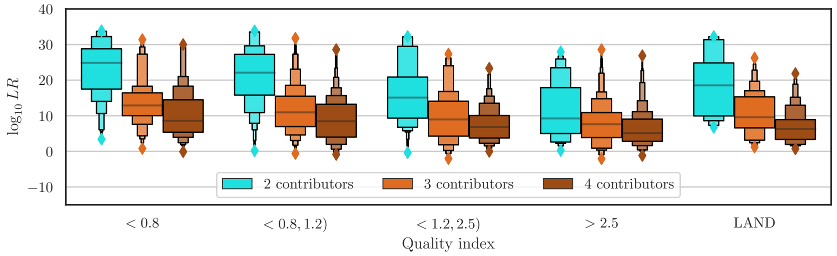


level of contribution indicate that HMC provides higher LRs than Euroformix in cases in which either the amount of template is low or the POI is a minor contributor to the mixture. This might be explained by the higher analytical threshold used by Euroformix, which is needed to avoid the presence of unmodelled double-backward stutters.

Finally, we test how $\log_{10}$ LR correlates with the quality index (QI) of the PROVEDIt mixtures. The QI is defined as the ratio between the concentrations of 80 and 214 base-pair fragments. When the concentration of 214 base-pair fragments is below detection level, the QI can not be defined and the mixture is instead categorized as "LAND" (large autosomal not detected) [2]. The QI can serve as a simple metric quantifying the level of sample degradation. The higher the QI, the lower the expected quality of the EPG peaks. As expected, the $\log_{10}$ LR are therefore lower for mixtures with higher QI (Figure 7). The median $\log_{10}$ LR for mixtures with QI less than 0.8 is 13.44 (7.941 ... 18.748), it is 12.23 (6.974 ... 18.086) for QIs between 0.8 and 1.2, 8.684 (4.819 ... 14.304) for QIs between 1.2 and 2.5, and 6.98 (3.153 ... 10.448) for QIs higher than 2.5. LAND mixtures, however, do not follow this trend. 24 out of the 25 LAND mixtures (138 out of 162 profiles) have had their quality degraded by the largest used volume of humic acid (35 µl). Humic acid is known to affect the large autosomal target in the Quantifiler kit that was used for the quantification of the mixtures [1]. However, humic acid does not seem to affect the other STR peaks too much. For example, one can compare the untreated mixture

```
A03_RD14-0003-40_41-1;4-M3I35-0.315GF-QLAND_01.15sec
```

with the mixture

```
G02_RD14-0003-40_41-1;4-M3a-0.315GF-Q0.6_07.15sec
```

treated with the highest volume of humic acid. Both mixtures contain the same amount of DNA material in the same proportions between the same contributors. The peak heights do not indicate significantly stronger degradation when humic acid is used.

The differences between results computed by HMC and STRmix™are larger when the QI is low (i.e. the quality of the mixture is high), as shown in Figure 8. We show this by analysing the standard deviation of the differences between the solutions, as the medians of the differences are all close to zero (e.g. 0.037 for QI less than 0.8 and -0.011 for QI greater than 2.5 in case of 2-contributor mixtures). Comparing HMC with STRmix™for 2-contributor mixtures, the standard deviation of the differences in the $\log_{10}$ LR is 1.241 for QI less than 0.8 and 0.836 for QI greater than 2.5. For 3-contributor mixtures, the standard deviations of the differences for the same QI classes are 1.514 and 0.482, respectively, and for 4-contributor mixtures they are 1.276 and 0.589. These results contradict the intuition, as one would expect larger differences when the information content of the mixtures is lower. In such a case the difference would be more likely caused by noise in the data. Comparing HMC with Euroformix

Figure 8: Distributions of the differences between the $\log_{10}$ LR obtained by different methods when true hypotheses are explored for mixtures with different PROVEDIt quality indices and different numbers of contributors.

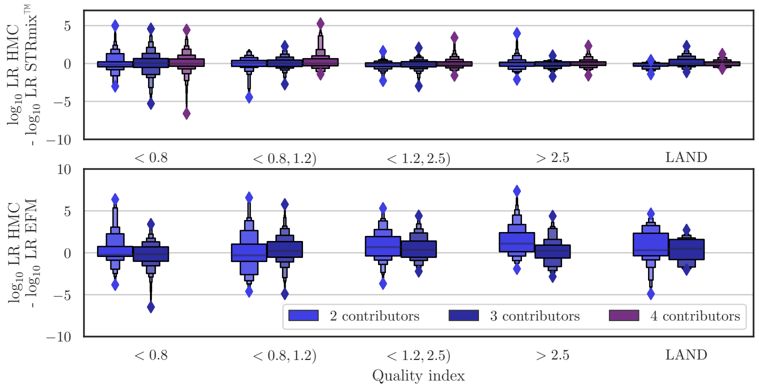


(EFM) shows standard deviations of the $\log_{10}$ LR differences of 1.822 and 1.83 for QI $< 0.8$ and $> 2.5$, respectively, for 2-contributor mixtures, and 1.512 and 1.508 for 3-contributor mixtures. But in this case, the medians of the differences display a trend. In case of 2-contributor mixtures the median is larger in the case of low quality samples (1.085) than in the case of high quality samples (-0.235).
