## Supplementary Material 2 for "Analysis of the Hamiltonian Monte Carlo genotyping algorithm on PROVEDIt mixtures including a novel precision benchmark"

—

—

### Analysis of the 4-contributor mixture with a large difference between the reported results

We provide an analysis of the scenario consisting of the mixture E05_RD14−0003−33_34_35_36 −1;4;4;1−M2a−0.63GF (E05), a hypothesis of the defendant with 4 unknown contributors, and the hypothesis of the prosecutor with 3 unknown contributors plus the POI "Contributor 33". HMC reports $\log_{10}$ LR = −0.0852, while STRmix™reports 6.5106. To understand the possible sources of this stark difference, we look for the loci that do not provide support for the hypothesis of the prosecutor. HMC suggests that there are two minor contributors. The estimated DNA proportions/weights for them are between 0.09 ... 0.16 and 0.02 ... 0.09, respectively. The second potential minor contributor matches well another true POI, "Contributor 36" with a corresponding sub-sub-source $\log_{10}$ LR of 9.5455. The first potential minor contributor has the highest sub-sub-source $\log_{10}$ LR for "Contributor 33", namely 0.5166. We therefore focus on Contributor 33.

The following loci had sub-sub-source LRs lower than 1 for Contributor 33:

- CSF1PO: sub-sub-source LR 0.218
- D10S1248: 0.855
- D12S391: 7.769e−12
- D16S539: 0.942
- D8S1179: 0.178
- vWA: 0.269

In the following, we focus on locus D12S391, which provided the strongest support for exclusion. The minor contributor's genotype is (18, 18.3). We note, however, that the peak 18.3 is not called (Figure 1). In our estimation of the posterior, the allelic product for this contributor (including multiplication over the estimated values of the amplification efficiency of D12S391) ranges from 480 to 2040 RFU. The analytical threshold is 60 RFU, so a dropout is highly unlikely, rendering the HMC result plausible on these input data.

For comparison, let us consider the alternative scenario in which peak 18.3 is called. We estimate the height of this peak to be around 1220 RFU from the graphical user interface of Genemapper® (Figure 1). Table 1 compares the resulting LRs with the previously reported ones. We observe a strongly increased $\log_{10}$ LR value from HMC upon including peak 18.3 in the input data. The ENFSI verbal equivalent of the evidence strength changes from *"no support for one of the propositions over the other"* without peak 18.3 to *"provides extremely strong support for the proposition of the prosecutor over the other"* with peak 18.3. At the same time, the LR values for the other POIs do not change significantly.

This example highlights once more how important it is for practitioners to understand the mixture analysis framework and to be aware of possible pitfalls. Software that is used in the analysis

Figure 1: A screenshot from GeneMapper® showing the D12S391 locus of the E05 mixture. Allele 18.3 has not been called, despite a visible shoulder.

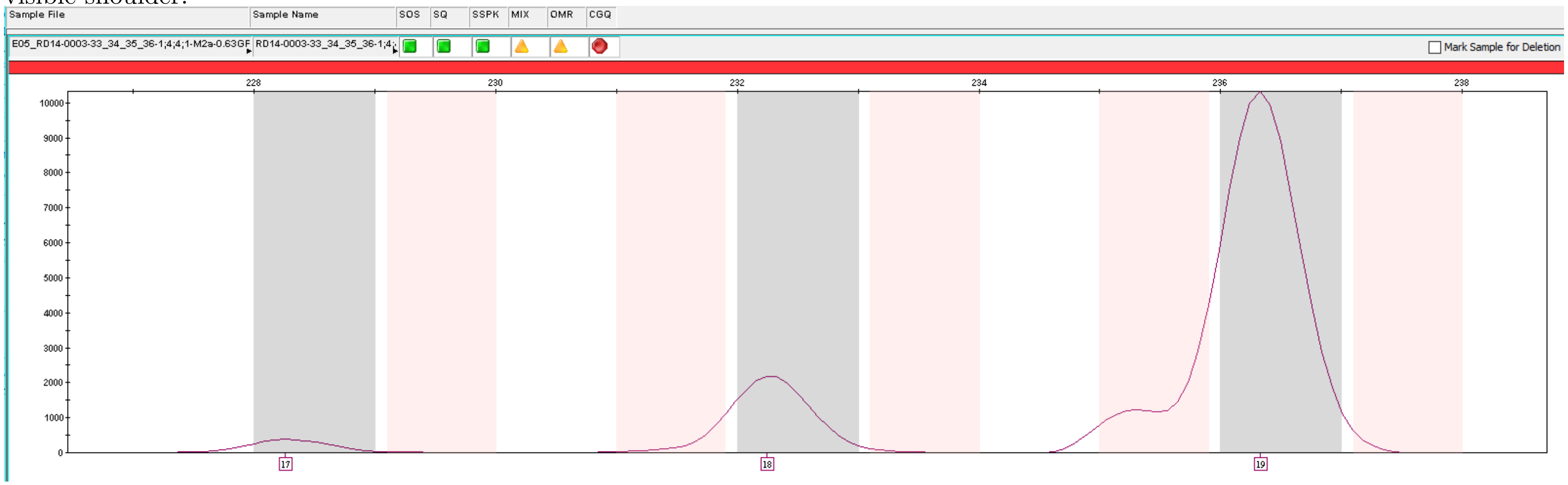


| Contributor | 18.3 called HMC | 18.3 not called HMC | 18.3 not called STRmix™ | 18.3 not called EFM 2.1.0 |
|---|---|---|---|---|
| 33 | **12.8367** | **-0.0852** | 6.5105 | 9.132 |
| 34 | 19.9202 | 19.7855 | 19.1173 | 18.6257 |
| 35 | 19.7403 | 19.6019 | 19.1399 | 19.1399 |
| 36 | 8.8806 | 8.9406 | 5.5832 | 5.5832 |
| C41C_Cauc | -45.767 | -45.501 | $-\infty$ | -2.496 |
| MT95124_AA | $-\infty$ | $-\infty$ | $-\infty$ | -3.969 |
| PT84215_AA | $-\infty$ | $-\infty$ | $-\infty$ | -4.878 |
| PT84223_AA | $-\infty$ | $-\infty$ | $-\infty$ | -5.01 |

Table 1: $\log_{10}$ LR for different POIs on the E05 mixture. $H_p$ assumes three unknown contributors plus the POI, and $H_d$ assumes four unknown contributors. Two scenarios are considered for HMC inference: with and without the 18.3 peak in locus D12S391 called. Contributor 33's genotype includes the previously not called allele 18.3. The Euroformix (EFM) results are taken from the NIST study [1].

process should provide easy to understand diagnostics in order to limit the need for sophisticated manual analysis. An improvement would be to include "possible shoulder peaks" in the output of the peak calling algorithm. If Genemapper® or its alternatives would report possible shoulder peaks and the corresponding certainty scores (the higher the more likely it is that such a shoulder is indeed a peak), the probabilistic genotyping model could take this information into account during the inference process.

An analyst taking care of the LR calculation should notice that the 18.3 peak has not really been dropped. It is present in the Genemapper software and simply was not resolved. A similar case had already been discussed by Russell et al. [2], where, unlike in the case described here, the computed LR was 0. That result was, however, likely caused by the STRmix™ behaviour discussed in Subsection 3.1 of the main text.
